# A numerical bias in honeybees: Numerousness is more salient than space and size non-numerical cues during quantity discrimination

**DOI:** 10.64898/2026.03.25.714149

**Authors:** Elena Kerjean, Scarlett R. Howard, Aurore Avarguès-Weber

## Abstract

Despite growing evidence that many animals can evaluate quantities, the ecological relevance of numerical cognition remains debated, particularly outside vertebrates. Would individuals still rely on numerousness if less computationally demanding cues, visual features extracted at the early stage of visual processing, were available to assess quantity? In primates, individuals show a numerical bias as they tend to rely on the number of items rather than non-numerical cues, such as total area, to categorize quantities. In this study, we trained free-flying honeybees to discriminate between two and four items in conditions where numerosity covaried with the total area and perimeter (Experiment Size) or the convex hull (Experiment Space) cues, mimicking ecological contexts. Transfer tests assessed which numerical or non-numerical cues were learned and preferentially used by the bees. Bees primarily relied on numerousness over these non-numerical cues. Individual analyses revealed two consistent strategies: a “numerical bias” strategy, in which bees encoded numerical information while ignoring non-numerical cues, and a “generalist” strategy, where bees flexibly switched between cues and favored non-numerical information when cues conflicted. We further reported improved discrimination when smaller quantities appeared on the left and larger ones on the right, consistent with an oriented mental number line. Together, these findings demonstrate a spontaneous numerical bias in honeybees and reveal that individuals within the same species can adopt distinct strategies when evaluating quantity. Our findings also suggest that distantly related taxa like bees and primates may have independently evolved comparable mechanisms for quantity evaluation.

## Introduction

Animals benefit from discriminating “more” from “less” in several situations (Bortot et al., 2021; Nieder, 2021), yet, the reasons they would rely on abstract numerousness rather than other perceptual cues remain an open question (Gebuis et al., 2016; Leibovich et al., 2017; Núñez, 2017). In the wild, numerousness almost always covaries with modality-dependent non-numerical cues of quantity, such as cumulated area, perimeter, or density, which are extracted early on in sensory neuronal pathways. Numerousness, by contrast, requires a higher level of neuronal integration and is therefore assumed to be more computationally demanding.

Numerousness defines the cognitive representation of the approximate “number” of elements within a set (i.e., its numerosity). Numerousness is an abstract concept of numerosity independent of the physical properties of the elements (Dos Santos, 2022; Núñez, 2017; Stevens, 1936). By definition, numerosity is a precise measure of a set’s cardinality, often expressed using numerals (e.g., “1”, “2”, “3”) and only accessible using counting-like mapping processes, while numerousness is an inherently approximate representation of the precise numerosity of a set. The ability to compute numerousness is thought to rely on an evolutionarily conserved Approximate Number System (ANS) (Cantlon & Brannon, 2006), described across vertebrates (see recent reviews in fish: Agrillo and Bisazza, 2018 and birds: Regolin et al., 2025) and potentially present beyond (honeybees: Howard et al., 2018, 2019b, fruit flies: Bengochea et al., 2023). Such shared capacity is also defined as “number sense” (Dantzig, 1931; Dehaene, 1997).

Many non-numerical cues of quantity could also inform the question “how much is there?”. Within each sensory modality, a plethora of non-numerical cues correlating with quantity can be described (for a recent review, see Shilat et al., 2024). Existing classifications, such as the intrinsic versus extrinsic distinction (DeWind et al., 2015; Piazza et al., 2004), have sometimes added confusion in the field (Shilat et al., 2024). To clarify terminology, we adopt here an intuitive, yet potentially incomplete, distinction between Size and Space non-numerical cues, consistent with recent visual stimulus-standardization programs (Guillaume et al., 2020; Zanon et al., 2022). Size cues relate to item dimensions (e.g., total area, total perimeter, item size, coverage), while Space cues depend on items’ spatial arrangement (e.g., convex hull, density, inter-distance, spatial frequency). Space cues are limited by Size cues, so that, for example, the convex hull area (e.g., the area of the smallest polygon surrounding all items in a set) cannot be larger than the total area of items (Guillaume et al., 2020). Although related, this framework does not fully overlap with the seminal model of DeWind et al. (2015), as in our work Size and Space relate to categories and not computational variables implemented within a model, as it is the case in DeWind et al. (2015).

Numerousness, Size, and Space cues are inherently intertwined and frequently co-vary in natural scenarios. Together, these magnitudes fall under the scope of quantity discrimination, which enables individuals to discriminate “more” from “less” using one or a combination of these cues. This ability is highly advantageous and has been documented across the entire animal realm. Many species spontaneously discriminate quantity in foraging contexts, including terrestrial mammals (Bánszegi et al., 2016; Cox & Montrose, 2016; Hauser et al., 2000; Irie-Sugimoto et al., 2009; Rivas-Blanco et al., 2020; Schaffer et al., 2025; Snyder et al., 2021; Uller & Lewis, 2009), marine mammals (Abramson et al., 2011, 2013), birds (Aın et al., 2009; Garland et al., 2012), non-avian reptiles (Cooper et al., 2024), amphibians (Krusche et al., 2010; Stancher et al., 2015), cartilaginous fish (Kreuter et al., 2021), and cephalopods (Yang & Chiao, 2016). However, quantity is not only a matter of food. Fish and flies, for example, preferentially approach larger groups of conspecifics, benefiting from dilution effects to avoid predation (Agrillo et al., 2008; Ferreira & Moita, 2020). Other species use quantity cues to assess intra- or inter-specific competition (mealworm beetles: Carazo et al., 2012; hyenas: Benson-Amram et al., 2011) or to increase their chances of finding suitable habitat (tree frogs: Lucon-Xiccato et al., 2018; dune snails: Bisazza and Gatto, 2021). Recent work even suggests that quantity discrimination may extend to plants (Guerra et al., 2025). Surprisingly, pea plants appear to orient their growth, maximizing the likelihood of contacting “more” rather than “less” vertical supports.

Despite the wealth of research demonstrating quantity discrimination in animals, it often remains unclear to what extent numerical and non-numerical cues drive behavior in natural contexts (Gatto et al., 2022; Messina et al., 2022). To investigate non-numerical cognition, most studies have attempted to isolate these dimensions from each other, for example, by equalizing non-numerical cues between stimuli differing in quantity (Piazza et al., 2004). One limitation to this approach is that a single pair of stimuli that vary in numerosity cannot perfectly eliminate all non-numerical cues at once. For example, if two dot arrays have, respectively, two and four dots, and we equalize the total area, then the two dots will have a larger diameter than the four dots. Several techniques have been proposed to minimize the influence of non-numerical cues (see recent methodological review Shilat et al., 2024). A widely used method mixes learning trials in which numerosity is congruent or incongruent with specific non-numerical cues, preventing those cues from being predictive of a given quantity in 100% of the learning trials (Bortot et al., 2019; Ditz & Nieder, 2016; Gebuis & Reynvoet, 2011; Zanon et al., 2025). Yet, this solution can only handle a restricted set of cues and cannot prevent animals from switching between cues from one trial to the next. To address this issue, a complementary approach explicitly models the relative contributions of numerical and non-numerical cues during quantity evaluation (Aulet & Lourenco, 2023; Cicchini et al., 2016; DeWind et al., 2015; Ferrigno et al., 2017; Starr et al., 2017). This method allows researchers to quantify the influence of both numerical and non-numerical cues during quantity evaluation. In humans these studies provide a line of evidence showing that numerousness is often more salient to individuals than non-numerical magnitudes during quantity evaluation (Cicchini et al., 2016; DeWind et al., 2015; Ferrigno et al., 2017; Starr et al., 2017).

Such preferred use of numerousness is known as the “numerical bias” (Aulet & Lourenco, 2023). Experiments explicitly assessing this relative cues bias often follow a stepwise protocol: (1) during training, numerosity and a non-numerical cue covary; (2) during testing, novel stimuli dissociate these dimensions: some arrays vary only in one dimension while others vary in the other, making individuals unable to rely on one or the other cue. This design reveals which cue individuals relied on during learning. Ferrigno et al. (Ferrigno et al., 2017) applied a version of this methodology to compare the saliency of numerousness and total area in adults from high- and low-numeracy cultures, children, and rhesus macaques. They used a two-dimensional stimulus matrix varying in either total area (8-22 *cm*^2^) or numerosity (8-22 dots). During training, only perfectly correlated stimuli were presented, and participants learned under an appetitive-aversive paradigm to classify “small” versus “large” arrays in two categories. In the testing phase, they started to dissociate the correlation between the two cues. The respective influence of each magnitude on the classification made by the subjects could then be modeled. The data shows that across all groups, individuals tended to classify stimuli according to numerousness rather than area, revealing a shared “numerical bias” across cultures, ages, and species. The bias was, however, stronger in humans compared to non-human primates, and within humans, in adults compared to children.

Beyond primates, our understanding of numerical bias remains limited for several reasons. First, quantity-discrimination tasks often rely on spontaneous preferences, which tie behavior to ecological context, making it difficult to determine whether observed preferences reflect adaptive constraints or relative cue saliency. For example, in a food choice assay, dogs spontaneously rely on total area rather than numerousness (Petrazzini & Wynne, 2016). Such strategies may be adaptive in a given context, therefore not necessarily reflecting the intrinsic saliency of total area in other conditions. Similar reasoning can be applied, for example, to the saliency of density cues in fish tendency to join larger shoals (Gómez-Laplaza & Gerlai, 2013) given the specific adaptive value of density regarding dilution effects (Wrona & Dixon, 1991). A second limitation arises when animals are trained with numerosity and non-numerical cues fully correlating but tested only controlling for the non-numerical dimensions during numerical transfer tests, but often without introducing a symmetrical non-numerical transfer test. In such a procedure, if animals succeed in the numerical transfer tests, it indicates that they have spontaneously encoded numerousness, however, it does not inform on the relative saliency of numerousness versus other non-numerical cues. Hence, it is impossible to fully conclude from these studies if a numerical bias exists in the species (see, for example, studies in chicks (Rugani et al., 2008, 2014)). By contrast, if animals fail in these numerical transfer tests, it rather suggests that they may not use numerousness as a spontaneous cue of quantity. Such absence of numerical bias has, for example, been reported in fish using visual and tactile quantity discrimination (Agrillo et al., 2010; Bisazza et al., 2014) and in cats using visual dot arrays (Pisa & Agrillo, 2009). Finally, the limited number of studies, mainly involving pigeons (Diaz and Wasserman, 2023, n.d.; Kubo, 2022), that have reported both numerical and non-numerical transfer tests after quantity discrimination training concluded that individuals encoded both numerical and non-numerical cues without numerical bias. Such studies have not yet been undertaken outside vertebrate species. Recent evidence in fruit flies, however, suggests a prominent role of non-numerical perceptual magnitudes (Bengochea et al., 2023).

Honeybees constitute a valuable model for exploring the existence of a “numerical bias”. Their brains contain fewer than one million neurons (Menzel & Giurfa, 2001), and their lineage diverged from primates more than 600 million years ago (Nieder, 2021). Yet, their surprisingly sophisticated “number sense” (Dantzig, 1931; Dehaene, 1997) includes abilities previously thought to be restricted to primates, such as representing zero Howard et al., 2018. Over the last decades, honeybees have indeed progressively become a major model for studying abstract rule learning outside the vertebrate clade (Avargues-Weber & Giurfa, 2015; Avarguès-Weber et al., 2011, 2012, 2014; Giurfa et al., 2001). Pioneered research demonstrated proto-counting abilities (Chittka & Geiger, 1995) and quantity generalization across visual stimuli (Dacke & Srinivasan, 2008). More recent studies using appetitive-aversive differential conditioning (Avarguès-Weber, de Brito Sanchez, et al., 2010) showed that honeybees can perform numerousness judgments of small (1-4) and large (5-8) numerosities while controlling for Size cues (Howard et al., 2019b) or for both Size and Space cues (Bortot et al., 2019). Beyond numerousness judgment, their ability to manipulate numerical information is unprecedented outside vertebrates: honeybees can perform elemental addition and subtraction (Howard et al., 2019a), learn to match symbols to quantities (Howard et al., 2019c), order small quantities, and extend this ordering to include the concept of zero, following typical signatures of the ANS (i.e., distance effect during discrimination between zero and progressively larger numerosities; Howard et al., 2018). These studies, however, raised debates on the potential alternative use of non-numerical cues by the bees, including total perimeter, convex hull area, and spatial frequency (MaBouDi et al., 2021; Shaki & Fischer, 2020). Given that the honeybee brain consists of fewer than one million neurons, one might indeed reasonably expect bees to favor non-numerical cues whenever these are available.

In this study, we assessed the preference of honeybees (*Apis mellifera*) to use numerical versus non-numerical magnitude cues in quantity discrimination. To do so, we trained bees to discriminate dot arrays in which numerosity covaried with selected non-numerical cues, adapting protocols used in number bias studies in primates and birds (Diaz & Wasserman, 2023; Ferrigno et al., 2017; Kubo, 2022). Free-flying foragers were trained in an appetitive-aversive quantity discrimination task (2 vs. 4). In the first experiment, items’ sizes were equalized so that numerousness and Size cues (i.e., total area and perimeter) covaried during training, allowing bees to rely either on numerical or non-numerical Size cues to solve the task. Space cues were partially controlled by homogenizing the convex hull, density, and mean inter-distance across sets of training stimuli. After training, bees were subjected to non-reinforced transfer tests and a conflict test. In the numerousness transfer test, Size cues were discarded by equalizing total area between the test stimuli differing in the number of items displayed. In the Size transfer test, the use of numerousness was prevented by homogenizing numerosity and assessing bees’ preference when stimuli presented similar total area to the rewarded and non-rewarded stimuli during training but displayed the same number of items (3 in each stimuli). Finally, in the conflict test, the correlation between cues was reversed from training: smaller numerousness being paired with larger Size cues and vice versa. A second experiment followed the same training and testing pattern, except that Space cues (i.e., convex hull) and numerosity co-varied, while Size cues were controlled.

If bees spontaneously encode numerousness we expected them to perform higher than chance level during numerical transfer tests in both experiments. If bees present a “numerical bias”, they additionally should perform better in numerical transfer tests rather than in Size and Space transfer tests. The conflict test enables us to measure the extent to which numerousness is evaluated independently from non-numerical quantity cues. Together, these experiments allowed us to assess whether numerousness spontaneously guides honeybees’ decisions and provided a direct test of whether a numerical bias exists outside the primate lineage.

## STAR METHODS

### EXPERIMENTAL MODEL AND STUDY PARTICIPANT DETAILS

Data for Experiment Size were collected from January to March 2024 at the Jock Marshall Reserve at Monash University’s Clayton campus (Melbourne, Australia). Sixteen free-flying forager honeybees (*Apis mellifera*) participated in the experiment and originated either from two managed hives or a nearby wild colony. Foragers belong to the non-reproductive female worker caste and occupy the final stage of the age-related division of labour (Johnson, 2010); they typically range from 20 to 42 days old before dying naturally (Seeley, 2009). Data for Experiment Space were collected from June to July 2024 at the Toulouse University research apiary (Toulouse, France), including approximately twenty-five hives, and similarly included sixteen forager honeybees. In both experiments, bees initially visited a gravity feeder providing a 5-20% sucrose solution (weight/weight; as in Howard et al. (2019a)). Concentration of sugar feeder needs to vary across the season to maintain an equivalent amount of recruitment (i.e., higher concentration early summer, lower concentration end of summer). Individuals were then offered a drop of 50% (w/w; 1.8 M) sucrose on a spoon and transferred to the experimental setup, where they were placed on a small Plexiglas platform at the entrance of the Y-maze. Bees that later returned autonomously established a stable foraging route and were individually marked on the thorax and/or abdomen with POSCA water-based paint for identification. Although only one bee was trained at a time, several individuals could be recruited early on to increase the likelihood that at least one forager reliably located the apparatus from the hive.

### METHOD DETAILS

#### Study design

To ask whether honeybees rely on numerousness itself or on the non-numerical cues that usually covary with numerosity, we trained free-flying foragers in two complementary experiments. In both experiments, bees learned to discriminate stimuli presenting two dots from ones presenting four dots according to an appetitive-aversive conditioning paradigm. We did not include more numerosities to limit the effect of ratio and magnitude on our results. In Experiment Size, all dots were of the same size, as a consequence, numerosity covaried with the total area and the total perimeter. Stimuli with four dots presented twice the total area and total perimeter of stimuli with two dots. We controlled for Space cues (convex hull, density, and inter-distance) throughout the training phase. In Experiment Space, the same logic was followed, but here fixing the mean inter-distance between items made numerosity co-vary with the convex hull and controlled for Size cues (total area, total perimeter, and item size). In both cases, bees could solve the task using numerical and/or non-numerical cues. Following 40 learning trials, bees started the testing phase, including phase test types: two learning tests presenting stimuli similar to those presented during the training phase but never seen by bees before, one numerical transfer test, one non-numerical transfer test (Size or Space dependent on the experiment), and one conflict test in which one stimulus presented the target numerosity but the non-numerical cue value of the unrewarded stimuli during training, while the other presented the value of the non-numerical cue of the rewarded stimuli but with the incorrect numerosity.

#### Apparatus

The Y-maze structure was constructed from gray Plexiglas material and enclosed with a transparent Plexiglas ceiling (Figure 1), following the design described by (Avarguès-Weber et al., 2011). Stimuli were fixed to the center of the 20 × 20 cm back wall of each arm (*W idth* × *Height* × *Length*, 20 × 20 × 40cm), positioned 15 cm from the centroid of the decision chamber. This placement resulted in a visual angle of 26° (Giurfa et al., 1996). Each back wall was covered by a white 80 *µm* Lowell laminate cover. In the center of each stimulus, a small hole allowed the insertion of a pipette from which bees could drink the conditioning solutions.

**Fig 1.**
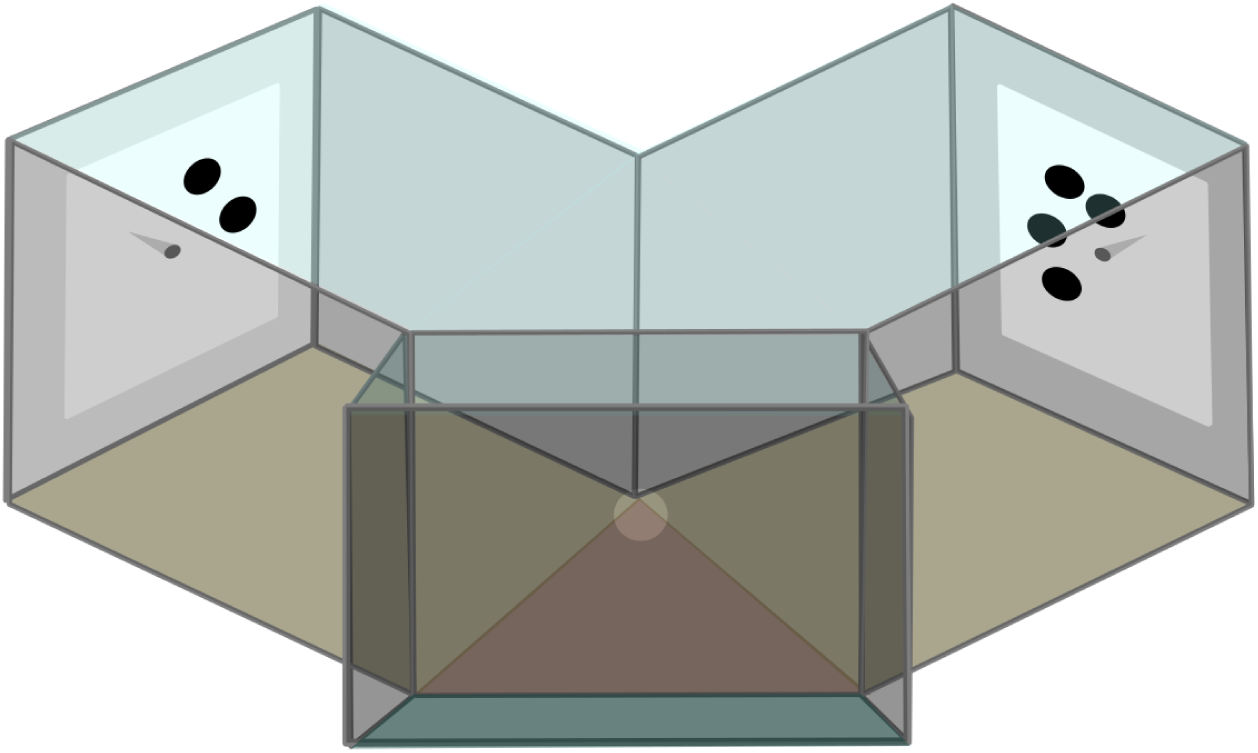
Schematic representation of the Y-maze. Bees entered the maze through a small slot created by slightly lifting the main entrance door. They first reached the entrance chamber (green), then passed through a central hole to access the decision chamber (orange). From there, individuals chose between the two arms of the maze (yellow), each displaying a stimulus and containing either a sucrose reward or a quinine solution at its center.

#### Pre-training procedure

During pre-training, each bee was familiarized with the Y-maze and guided toward the location of the sucrose reward. A small grey Plexiglas platform holding *ad libitum* 50% sucrose solution first attracted the bee to the entrance chamber. While the bee was feeding, the platform was moved stepwise through the maze—into the entrance, then gradually toward the central decision hole, and then the bee was guided to the transparent micropipette tip at the center of each back wall. On the following foraging bouts, the bee had to find the tip independently and was then trained to locate the pipette in the opposite arm. The training phase began once the bee consistently flew into both arms. This pre-training stage typically lasted 30-90 minutes.

#### Training and testing procedure

During training, each bee completed 40 consecutive trials according to a spaced training protocol (see Chapter 1), meaning bees made only one choice per foraging bout, approximately every 5 minutes, before returning to the hive and coming back for the next trial (Figure 2). A trial began when the bee entered the maze, passed through the central hole, and approached one of the two stimuli presented at the end of the arms. A choice was recorded when the bee came within 1 cm of a pipette. If the bee chose correctly, she found a pipette providing *ad libitum* 50% sucrose solution; if she made an incorrect choice, she found a pipette filled with a 60 mM quinine “bitter tasting” solution acting as an aversive outcome (Avarguès-Weber, de Brito Sanchez, et al., 2010). These solutions were perceptually similar and could not be discriminated without direct contact of the antennae, the tongue, or a leg. In each experiment, one group (N = 8) was trained to choose stimuli with four dots (CS+) and to avoid stimuli presenting two dots (CS-), the other group was trained to choose stimuli with two dots and avoid those with four dots. Between trials (2-10 min), stimuli were removed, cleaned with 20% ethanol (as in Howard et al. (2019a)), and replaced with novel stimuli. The pipettes were refilled with the corresponding solution. The sequence of stimulus pairs followed a pseudo-random sequence similar for all bees, and no pair appeared more than twice consecutively. The rewarded side was also pseudo-randomized, with no side repeated more than twice in a row. When a bee developed a strong side bias, meaning more than four consecutive choices on the same side irrespective of the reward, the experimenter could adjust the rewarded side. No adjustments were made after trial 30 to ensure comparable final training between bees. Training typically lasted 2.5-4 h.

**Fig 2.**
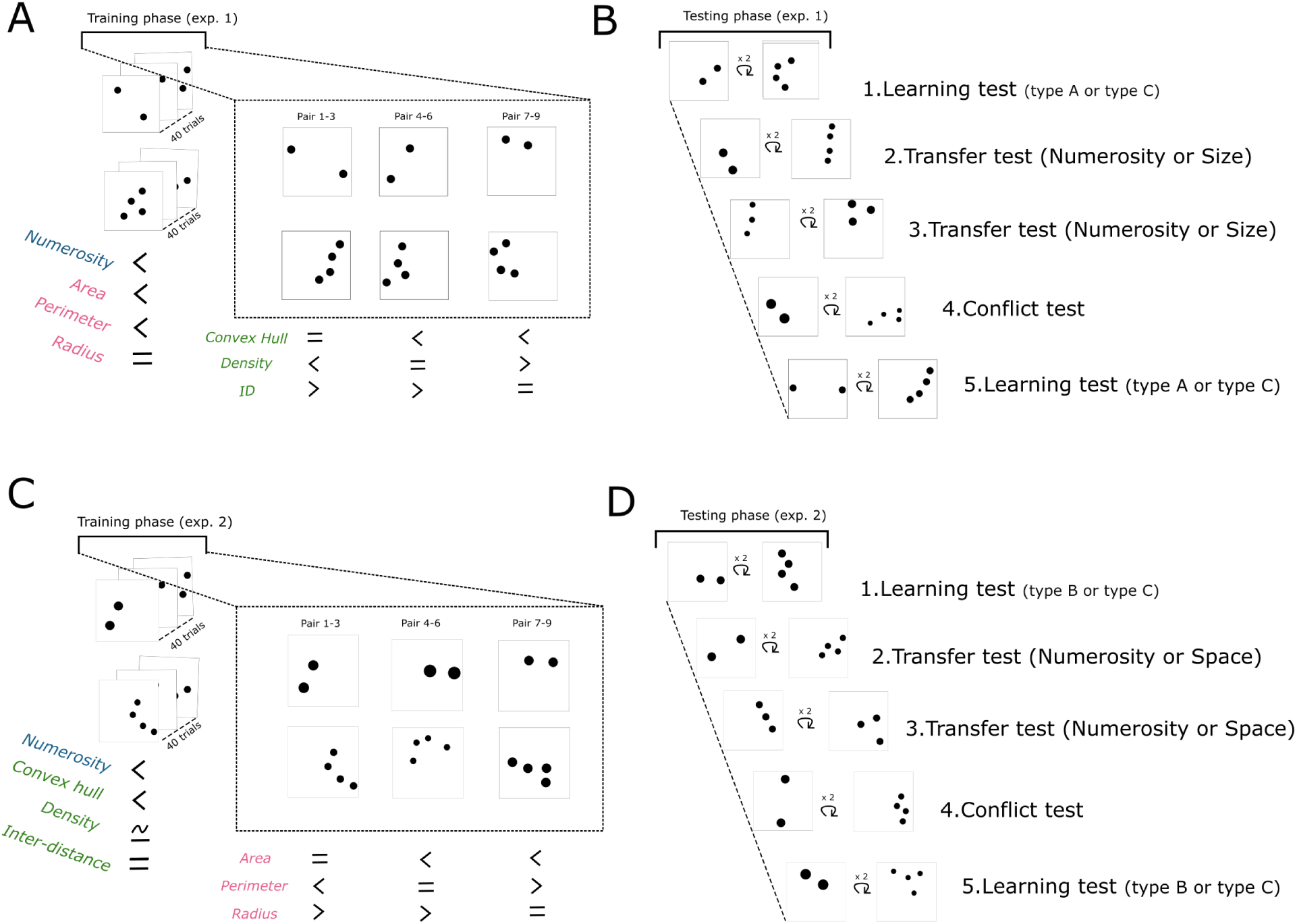
Protocol used in training and testing phases: A) Training phase protocol of Experiment Size. Bees are trained to discriminate between a set with two dots and a set of stimuli with four dots during 40 trials. Numerosity, total area, and total perimeter covary. Space cues are control over the training periods. Pairs A have a similar convex hull, pairs B have a similar density, and pairs C have a similar inter-distance. B) Testing phase protocol of Experiment Size. Five different test types are performed, each test is presented twice. The first test and last test are Learning Test A or C (randomized). The second and third tests are transfer tests, numerical or Size transfer (randomized). The fourth test is always the conflict test. C) Training phase protocol of Experiment Space. D) Testing phase protocol of Experiment Space.

The testing phase consisted of five test types, each comprising two unreinforced test rounds. In each round, the bee remained enclosed for 45 s in the maze with the test stimuli displayed. Novel pipettes were used, and the conditioning solutions were replaced with water as a neutral outcome. A test choice was recorded when the bee attempted to taste the water with its proboscis or antennae. Bees making fewer than 10 choices were allowed an additional 30 s. In the second round, we switched the side of the two stimuli, ensuring each test stimulus appeared once on each arm, and no side bias could account for our results. Between test rounds, bees received two reinforced refresher trials to maintain motivation and prevent extinction effects. Once both test rounds were completed, bees underwent three refresher trials before proceeding to the next test. The first and last test sessions were always learning tests, used to confirm that bees retained the trained rule throughout the testing phase. After the initial learning test, bees received one numerical transfer test (TN) and one non-numerical transfer test (Experiment Size: TS; Experiment Space: TSp), with the order counterbalanced across individuals. The third test was always the conflict test to minimize potential confusion for the bees during the transfer tests. Testing typically lasted 1.5-3 h.

#### Stimuli

Stimuli consisted of black dots on 15 × 15 cm white squares and were covered with 80 *µm* Lowell laminate. Nine pairs of stimuli were used during the training phase of each experiment. Each of the pairs consisted of a stimulus with four dots and another with two dots. Stimuli were generated using the open-source Matlab program GeNeSiS Zanon et al., 2022. For each stimulus, we recorded eight variables: numerosity, item size (IT), total surface (TA), total perimeter (TP), density (D), convex hull area (CH), mean inter-distance between stimuli (ID) and the total power without DC component (Table 1). Spatial frequency can be thought of as the rate of repetition of a pattern per degree of the observer’s visual field. It is equivalent to the number of repeating elements per unit of distance and is computed via a Fast Fourier Transform (FFT) (Cochran et al., 1967). The power spectrum derived from the FFT reflects the contribution of the different frequency components. The total power defines a single metric that sums all frequency contributions (i.e., the area under the power spectrum curve). It provides a single value enabling comparison between stimuli of similar pixel size and is commonly used in animal cognition studies (Adam et al., 2024; Adriano et al., 2021; Howard et al., 2022; Potrich et al., 2022). The first component of the Fourier transform corresponds to the luminance of the image (i.e., the amount of black pixels) and is often referred to as the DC component. To isolate spatial structure, we report the “total power” without the DC component, computed from (Zanon et al., 2022).

**Table 1.**
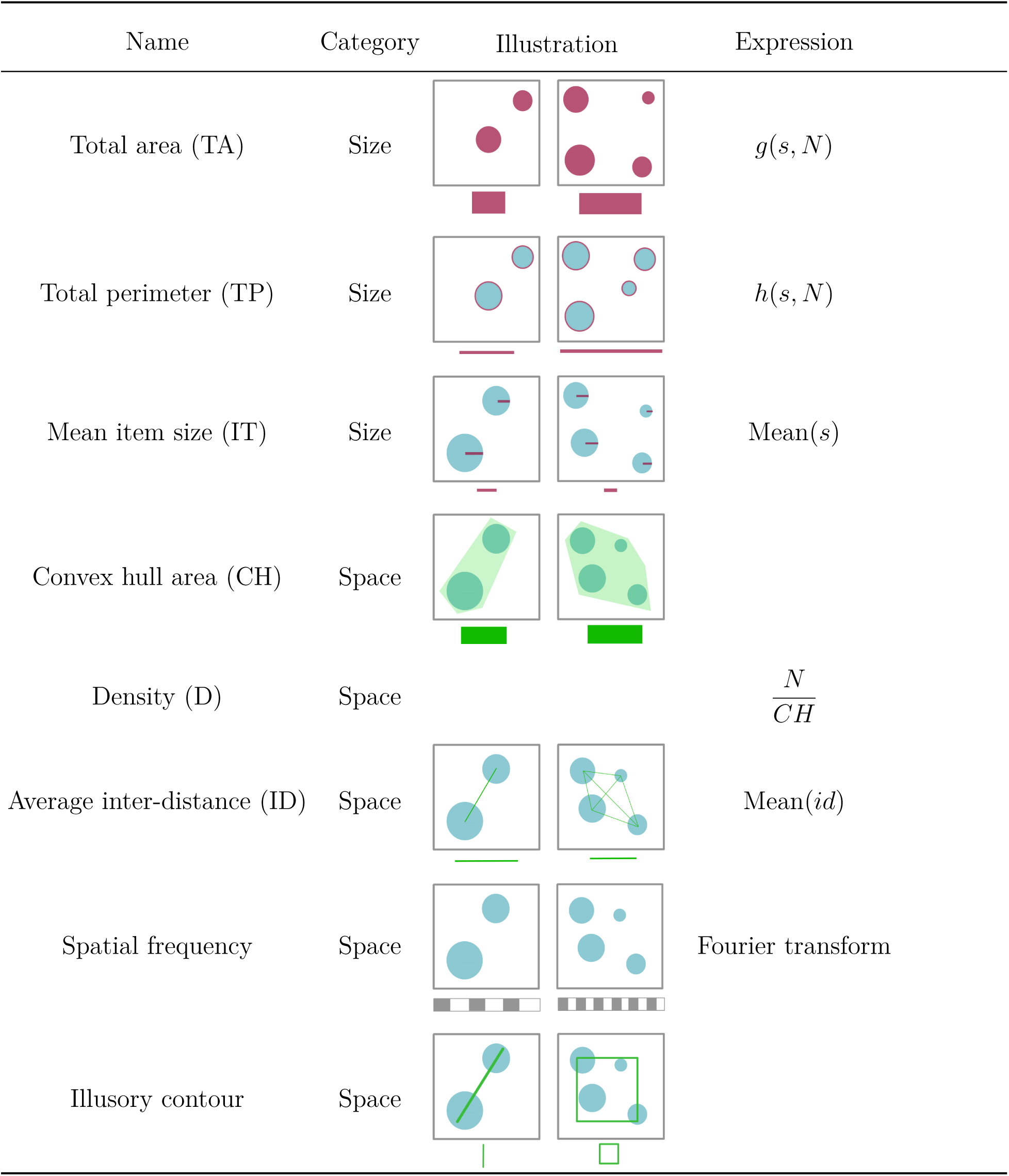
Description of non-numerical cues of quantity. Total area. (TA) represents the summed area of all items. **Total perimeter** (TP) represents the summed perimeter of all items. **Mean item size** corresponds to the mean size of all items. **Convex hull** (CH) defines the area of the smallest polygon that surrounds all items. **Density** (D) is the number of items divided by the convex hull area. **Average inter-distance** (ID) is the mean pairwise distance between items. **Spatial frequency** describes the rate of repetition of patterns or textures per degree of the observer’s visual field. **Illusory contour** refers to high-probability spatial arrangements of a small number of items that evoke a global shape. g: function, h: function, id: vector of all inter-distances, N : numerosity, s: vector of all item sizes.

### Experiment Size: Size vs. Numerousness

During training, individual item size (IT) was fixed at 0.9 cm, ensuring that total area (TA) and total perimeter (TP) covaried perfectly with numerosity and were both fully predictive of the reward. Each dot had a diameter of 1.8 cm, within the spatial acuity of a honeybee at a distance of 15 cm (Srinivasan & Lehrer, 1988). TA was fixed at either 5 cm^2^ (2 items) or 10 cm^2^ (4 items), and TP was fixed at either 11.2 cm (2 items) or 22.4 cm (4 items), both presenting a ratio similar to numerosity (r = 2; detailed stimulus values are provided in Table 2).

**Table 2.** Stimuli characteristics used during the training and testing phase of Experiment Size. Value of Numerosity, Size and Space cues of quantity in training and testing stimuli. In **bold**, are values covarying during the training phase. Ratios are expressed as the division of the value of the stimulus with the larger numerosity (or TA during TS) by the value of the stimulus with the smaller numerosity. Each value is the mean value of all similar stimuli. CH: convex hull, CNS: conflict test, D: density, ID: mean inter-distance, IT: item size, LT_A: learning test category A, LT_C: learning test category C, Num.: Numerosity, Power: power spectrum derived from Fourier transform without the DC component (freq = 0), TA: total surface, TN: Numerical transfer, TP: total perimeter, TS: Size transfer.

| Exp. 1<br>Size | Type | Num. |  | IT |  | TA |  | TP |  | D |  | CH |  | ID |  | Power |  |
| --- | --- | --- | --- | --- | --- | --- | --- | --- | --- | --- | --- | --- | --- | --- | --- | --- | --- |
|  |  | pair | ratio | pair | ratio | pair | ratio | pair | ratio | pair | ratio | pair | ratio | pair | ratio | pair | ratio |
| Training | A | 2 | 2 | 0.9 | 1 | 5 | 2 | 11.2 | 2 | 0.08 | 2 | 25.0 | 1 | 12.6 | 0.4 | 61.11 | 1.75 |
|  |  | 4 |  | 0.9 |  | 10 |  | 22.4 |  | 0.16 |  | 25.0 |  | 5.2 |  | 107.22 |  |
|  | B | 2 | 2 | 0.9 | 1 | 5 | 2 | 11.2 | 2 | 0.12 | 1 | 16.7 | 2.0 | 7.9 | 0.6 | 58.82 | 1.82 |
|  |  | 4 |  | 0.9 |  | 10 |  | 22.4 |  | 0.12 |  | 33.3 |  | 5.0 |  | 107.36 |  |
|  | C | 2 | 2 | 0.9 | 1 | 5 | 2 | 11.2 | 2 | 0.18 | 0.7 | 11.4 | 2.9 | 5.0 | 1 | 51.26 | 1.91 |
|  |  | 4 |  | 0.9 |  | 10 |  | 22.4 |  | 0.12 |  | 33.0 |  | 5.0 |  | 97.88 |  |
|  | Mean | 2 | 2 | 0.9 | 1 | 5 | 2 | 11.2 | 2 | 0.13 | 1 | 17.7 | 1.7 | 8.5 | 0.6 | 57.06 | 1.83 |
|  |  | 4 |  | 0.9 |  | 10 |  | 22.4 |  | 0.13 |  | 30.4 |  | 5.1 |  | 104.15 |  |
| Testing | LT_A | 2 | 2 | 0.9 | 1 | 5 | 2 | 11.2 | 2 | 0.08 | 2 | 25.0 | 1 | 12.6 | 0.4 | 57.11 | 1.99 |
|  |  | 4 |  | 0.9 |  | 10 |  | 22.4 |  | 0.16 |  | 25.0 |  | 5.2 |  | 113.48 |  |
|  | LT_C | 2 | 2 | 0.9 | 1 | 5 | 2 | 11.2 | 2 | 0.18 | 0.7 | 11.4 | 2.9 | 5.0 | 1 | 59.67 | 1.34 |
|  |  | 4 |  | 0.9 |  | 10 |  | 22.4 |  | 0.12 |  | 33.0 |  | 5.0 |  | 80.00 |  |
|  | TN | 2 | 2 | 1.1 | 0.7 | 7.5 | 1 | 13.7 | 1.4 | 0.14 | 1.1 | 14.7 | 1.8 | 5.0 | 1 | 92.12 | 0.66 |
|  |  | 4 |  | 0.8 |  | 7.5 |  | 19.4 |  | 0.15 |  | 26.4 |  | 5.0 |  | 60.76 |  |
|  | TS | 3 | 1 | 0.7 | 1.42 | 5 | 2 | 13.7 | 1.4 | 0.15 | 0.9 | 19.7 | 1.1 | 5.0 | 1 | 43.18 | 2.30 |
|  |  | 3 |  | 1.0 |  | 10 |  | 19.4 |  | 0.14 |  | 21.0 |  | 5.0 |  | 99.52 |  |
|  | CNS | 2 | 2 | 1.3 | 0.46 | 10 | 0.5 | 15.9 | 1 | 0.11 | 1.5 | 17.6 | 1.3 | 5.0 | 1 | 119.30 | 0.33 |
|  |  | 4 |  | 0.6 |  | 5 |  | 15.9 |  | 0.17 |  | 23.6 |  | 5.0 |  | 38.92 |  |

To limit the use of Space non-numerical cues, three categories of training stimuli were created, each equalizing one Space cue. Category A (pairs 1-3) equalized convex hull (CH), Category B (pairs 4-6) equalized density (D), and Category C (pairs 7-9) equalized mean inter-distance (ID). Across the full training set, mean cue ratios indicated that density was only weakly predictive of the reward (r = 1.22), whereas CH (r = 1.96) and ID (r = 0.68) remained predictive overall. We additionally computed the ratio of the total power for each type of pair (A, B, or C), which covaried with numerosity and Size during training (r = 1.83). In five of the nine training pairs, the four-dot pattern formed two clusters of two dots, whereas in the remaining four pairs, four dots formed a line of three plus one eccentric dot (Training type A example; Table 2). Finally, to reduce the risk that bees relied on simple positional heuristics, the two-dot stimuli were placed across the entire stimulus space (e.g., avoiding systematic placement of dots in the same corner). To evaluate their potential influence, two learning tests were included. Learning Test A (LT-A) used stimuli designed following Category A (CH equalized, IT predictive). Learning Test C (LT-C) used stimuli designed following Category C (ID equalized, CH predictive). In LT_C test the total power was less predictive (r = 1.34) than in LT_A (r = 1.99).

In the numerical transfer test (TN), the stimuli were homogenized for total area to the mean total area of training stimuli (7.5 *cm*^2^). Numerosity was unchanged. Here, the total power was incongruent with numerosity (r = 0.66). However, total perimeter (r = 1.4), convex hull (r = 1.8), and item size (r = 0.7) were congruent with numerosity. Their respective influence can be evaluated when integrating the results from the five different tests of both experiments (see Discussion). In the Size transfer test (TS), the numerosity of the stimuli was homogenized to the mean numerosity used during the training (3 dots), while total area was unchanged. Additionally, total perimeter (r =1.4), item size (r = 1.42), density (r = 0.93), convex hull (r = 1.07) and the total power (r = 2.30) were either partially or largely predictive of the total area. In the conflict numerosity size test (CNS), total area and numerosity of the stimuli were reversed compared to those used during the training (2 dots: TA = 5 *cm*^2^; 4 dots: TA = 10 *cm*^2^). The stimuli with 2 dots had a total area two times bigger than the one with 4 dots. The total power was also incongruent with numerosity in this test (r = 0.33). The transfer and conflict test had stimuli with similar inter-distance, like LT-C, this configuration enabled a larger diversity of dot positions on the stimuli. Two different sets of testing stimuli were used (1 and 2) with different positions of dots differing in illusory shape. Both four-dot numerical transfer stimuli formed a line to avoid any shape influence during this test.

### Experiment Space: Space vs. Numerousness

During training, the mean inter-item distance (ID) was fixed at 5.15 cm so that convex hull (CH) covaried reliably with numerosity. Because CH is jointly determined by spacing and item size, its exact value varied across stimuli, but the resulting ratios remained equal to those of numerosity (range: 1.4-3.1; mean r = 2). Density also changed across stimuli but was overall uninformative (overall r = 1), occasionally being mildly congruent (r = 1.3) or incongruent (r = 0.7) with numerosity. To limit the use of Size cues, three categories of training stimuli were created, each equalizing a different Size dimension: Category A (pairs 1-3) equalized total area, Category B (pairs 4-6) equalized total perimeter, Category C (pairs 7-9) equalized item size. Across the training set, density remained globally uninformative (r_D = 1.0), whereas TP (r_TP = 1.4) and IT (r_IT = 0.7) showed moderate predictive value. The mean predictability of the total power was uninformative (r = 1), however, in the stimuli of category C, it was congruent with numerosity and convex hull (r = 1.99).

As in Experiment Size, two learning tests were used to assess the potential influence of these cues: Learning Test B (LT-B; TP equalized, IT predictive, total power incongruent) and Learning Test C (LT-C; IT equalized, TP predictive, total power congruent). In the numerical transfer (TN), the convex hull was homogenized to 22.5 cm^2^, while numerosity remained unchanged. Under this configuration, density (r = 2), total perimeter (r = 1.4), item size (r = 0.7), and inter-item distance (r = 0.4) all varied consistently with numerosity, while the total power was incongruent (r = 0.7). In the Space transfer test (TSp), numerosity was homogenized to the mean trained value (3 dots), while CH values matched those encountered during training (15 vs. 30 cm^2^). Density (r = 0.5) and inter-item distance (r = 1.33) remained partially predictive, but not the total power (r = 1.1). In the conflict test (CNS), CH and numerosity were intentionally incongruent (2 dots: CH = 28 cm^2^; 4 dots: CH = 19 cm^2^). Owing to the geometric constraints of dot placements, CH could not be matched perfectly, but the resulting 1.5-fold still matched the congruency effect. In this test, the total power was incongruent with numerosity. TA was similar across transfer and conflict stimuli. Two sets of testing stimuli (Set 1 and Set 2) were used, differing in dot positions.

### QUANTIFICATION AND STATISTICAL ANALYSIS

#### General analysis procedure

Statistical models (Table S1) were assigned an identifier in the form x.y., where x denotes the experiment and y denotes the sequential order of analyses within each stage. All analyses were conducted in R (v4.2.3) via RStudio (v2023.03.0+386). All analyses were performed using generalized linear mixed models (GLMMs) with a binomial error distribution and logit link function, implemented in *lme4* (v1.1-32). Bee identity was included as a random intercept to account for repeated measures. For each model, the significance of predictors was assessed by comparing a full model to reduced models in which predictors were removed sequentially, using likelihood ratio tests (LRT) via the *drop1* function with *test = “Chisq”*. Where applicable, comparisons to chance level were performed using the *emmeans* function from the *emmeans* package (v1.2.2), tested against a null of 0 (logit scale) or 0.5 (response scale). The significance threshold was *α* = 0.05, and marginal trends were discussed when *α* ă 0.1. When multiple comparisons were required, p-values were adjusted using the Holm procedure. Post-hoc analyses of estimated marginal means (EMMs) were conducted using the *pairs*, *summary*, and *test* functions. For linear or categorical effects, we reported average marginal effects (AMEs), computed with the *margins* package (v0.3.26). Estimated marginal means (EMMs) were obtained using *emmeans*. When reporting results for categorical variables in the form A-B, the first level (A) corresponds to the model intercept.

Predictor selection was based on comparing all possible combinations of relevant predictors using the corrected Akaike Information Criterion (AICc), implemented via the *dredge* function from the MuMIn package (v1.47.5). Full models included biologically justified simple effects and two-way interactions. Simple effects selected through this procedure were retained regardless of statistical significance to limit omitted-variable bias. Biologically relevant two-way interactions were retained only when significant at *α* = 0.01. Each selected model underwent a three-step diagnostic evaluation. First, we checked for overdispersion using *check_overdispersion* from the *performance* package (v0.10.3). Second, we assessed multicollinearity by computing the variance inflation factor (VIF) for each fixed effect using *vif* from the *car* package (v3.1.2). Third, we examined model residuals by plotting residual deviance against predicted values and by inspecting binned residuals versus predicted values to ensure the absence of systematic patterns (e.g., linear increase or decrease, U-shape), following McCullagh and Nelder (1989).

Final models were compared to their corresponding null models (intercept + random effect) using the *anova* function with *test = “LRT”*, and results are reported in Table S1.

During the training phase, four predictors were considered: *trial* (learning trial 1-40); *type* (A, B, C), which specified the non-numerical control applied to the stimuli (*space_type* in Experiment Size; *size_type* in Experiment Space); *group*, indicating whether bees were trained to choose two dots (2:CS+) or four dots (4:CS+) stimuli; and *side*, specifying the Y-maze arm on which the correct stimulus appeared (L or R).

During the testing phase, predictors varied depending on the analysis. *test* denoted the type of test when multiple test types were modeled together; *group* indicated the trained rewarded numerosity (2:CS+ or 4:CS+); *test_order_all* specified the position of the test within the testing sequence (1-5); and *testing_set* denoted the specific stimulus pair seen by each bee.

#### Experiment Size

##### Dataset and final models

In total, the dataset included 16 individual bees, which completed the entire training phase (n = 640) and the five different test sessions (n = 3357). All final models can be found in Table S1. Here we detail the predictors evaluated for inclusion in the final model during the model selection process and the statistical coherence behind the each analysis.

##### Model 1.0: Training analysis

To analyze learning across training, we built a model (model 1.0; Table S1) testing the inclusion of : *trial* × *space_type* and *group* × *side*. Each interaction also included in dredge also tests for the simple effects alone. The response variable was the binary outcome of each learning trial (0 = incorrect, 1 = correct).

##### Models 1.1-1.3: Test analyses

For the testing phase, the response variable was again the binary choice on each test choice (0 = incorrect, 1 = correct). Because each test type could be differentially influenced by the predictors, we fitted three independent models: the two learning tests (model 1.1; LT_A and LT_C), the two transfer tests (model 1.2; TN and TS), and the conflict test (model 1.3; CNS). For models 1.1 and 1.2, we tested the inclusion of the following predictors: *test_order_all*, *group* x *side* x *test*, and *testing_set* x *test*. Interpretation focused on predictors involving *test*, while remaining predictors were retained as controls to avoid omitted-variable bias and ensure accurate inference. For the conflict test (model 1.3), no combination of predictors produced a satisfactory model fit (e.g., residual patterns in binned plots). We therefore retained a simplified model containing only the intercept and the random effect, and we interpreted conflict performance primarily through the clustering analysis, which yielded a better fitting model.

##### Model 1.4: Cluster and Mental Number Line analysis

To investigate whether distinct subgroups of individuals were present, we performed a clustering analysis based on bees’ performance in the two transfer tests and the conflict test. Hierarchical clustering was performed using the `eclust` function (*FUNcluster* = `hclust`, *hc_metric* = `pearson`). This approach assessed whether subsets of bees exhibited similar response patterns across tests. We evaluated solutions with *k* = 2 and *k* = 3 clusters. Only the two-cluster solution produced statistically meaningful separation; thus, we used the *k* = 2 solution to define a categorical predictor, *cluster*. In model 1.4, we examined the effects of *test* × *cluster* and *group* × *side* on bees’ choices (0 = incorrect, 1 = correct) across the entire testing dataset. Learning tests were included to determine whether clusters also explained variation in learning-test performance. We additionally compared corrected AIC values of models in which *cluster* was replaced by either *testing_set* or *group*. The model including *cluster* provided the best fit, confirming that this variable captured performance differences not explained by experimental grouping. We analyzed the Mental Number Line (MNL) effect (*group* × *side*) within this general model because the three-way interaction *test* × *group* × *side* was not significant.

##### Model 1.5: From spatial heuristics to rule learning

We additionally investigated *how* bees progressively discovered the visual discrimination rule in the Y-maze. We modeled the side chosen on trial *t* as a function of three predictors: the side chosen on the previous trial Choice_side(*t*–1), the side that yielded sucrose on the previous trial Reward (*t*-1), and the side on which the correct stimulus appeared on the current trial Correct (*t*). We analyzed the influence of these variables across training blocks. Inspection of binned residual plots indicated that residuals were not consistently centered around zero, suggesting a potential omitted variable. To limit the complexity of this exploratory model, we decided to limit the evaluation of additional predictors.

#### Experiment Space

The dataset included 16 individual bees, which completed the entire training phase (n = 640) and the five different test sessions (n = 2734). We conducted analyses parallel to those described for Experiment Size.

## RESOURCE AVAILABILITY

### Lead Contact

Further information and requests for resources should be directed to and will be fulfilled by the lead contact, Elena Kerjean.

### Materials Availability

Stimuli used during this experiment will be deposited at the time of publication jointly with the code and data.

### Data and Code Availability

- Behavioral data will be deposited at the time of publication and are publicly available as of the date of publication. Accession numbers are listed in the key resources table.
- All codes used for statistical analysis will be deposited at the time of publication and are publicly available as of the date of publication.
- Any additional information required to reanalyze the data reported in this paper is available from the lead contact upon request.

### Results Bees learned the quantity discrimination task

In Experiment Size, the two-dotted and four-dotted stimuli differed not only in numerosity but also in total area and perimeter. Bees (*N* = 16) could therefore rely on any of these cues to solve the task. Performance significantly increased across learning trials (Figure 3 A,B; trial: *AME* = 0.008, *p* ă 0.001). By the final trial, bees showed above chance (i.e, > 50%) performance (predicted mean at trial 40: 0.71, CI = [0.64, 0.77]). Learning did not differ between Space-controlled stimulus types (*space*_*type* : *p* = 0.568). After completing the 40 reinforced trials, bees were tested with two unreinforced learning tests that controlled different spatial cues. In LT_A bees could not use the convex hull, but inter-item distance was covarying with numerosity and Size, while in LT_C bees could not use inter-distance but the convex hull was covarying with numerosity and Size. LT_A and LT_C order in the testing phase was randomized. They performed significantly above chance in both tests (Figure 3 C; LT_A: mean = 0.63, CI = [0.59, 0.68], *p* < 0.001; LT_C: mean = 0.57, CI = [0.53, 0.61], *p* = 0.002), and performance in LT_A exceeded that in LT_C (*p* = 0.011).

**Fig 3.**
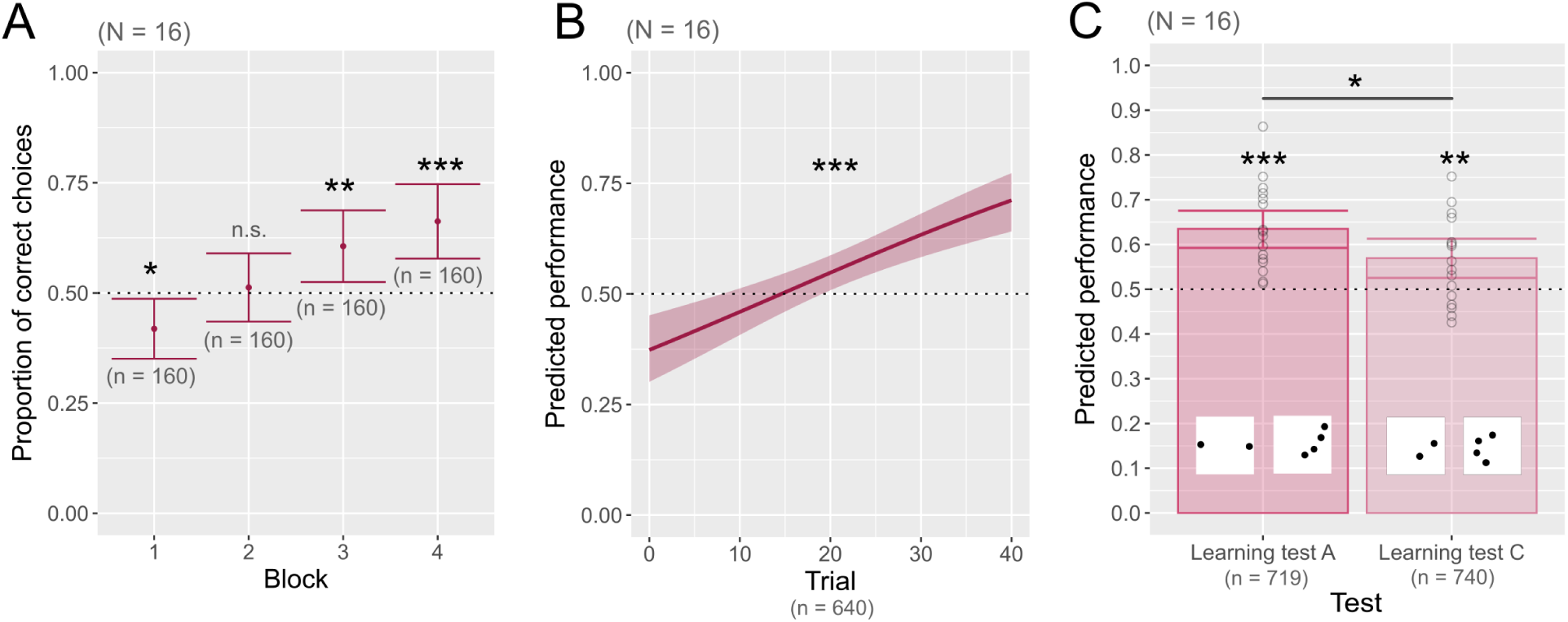
Experiment Size: Bees learned the quantity discrimination task. A) Mean proportion of correct choices across training blocks (Number of individuals: N = 16; Number of observations: n = 640). Each point represents the mean performance for a block of 10 trials. Error bars denote 95% CIs. Values are computed from raw data. B) Predicted probability of making a correct choice across trials (N = 16, n = 640). The solid line represents the prediction from model 1.0. Shaded areas indicate 95% CIs. C) Predicted performance in the two learning tests (N = 16; LT_A: n = 719; LT_C: n = 740). LT_A corresponds to the test in which the convex hull was controlled, and LT_C to the test in which inter-distance was controlled. Error bars denote 95% CIs. Data and significance are computed from model 1.1. Each dot represents the performance of a bee in one test bout (45 sec.) computed from raw data. The dotted horizontal line shows chance level (0.5). (***): p ≤ 0.001, (**): p ≤ 0.01, (*): p ≤ 0.05, (■): p ≤ 0.1, (n.s.): p > 0.1. N: number of individuals; n: number of observations.

In Experiment Space, the two-dotted and four-dotted stimuli differed systematically in numerosity and convex hull area. Bees (*N* = 16) again increased significantly in performance across trials (Figure 4 A,B; Figure S1 B; trial: *AME* = 0.005, *p* = 0.002). Predicted probability of making a correct choice at the final trial was 0.65 (CI = [0.58, 0.72], *p* < 0.001). Performance differed across the three Size-controlled stimulus types (Figure S2 A,B; *size*_*type*: *p* = 0.033): bees performed best when all dots had identical size (type C), a configuration in which total area and total perimeter were jointly congruent with numerosity and convex hull. Performance for type C (controlled item size) was significantly higher than for type B (controlled perimeter) (C-B: *AME* = 0.12, *p* = 0.046) and tended to exceed type A (controlled area) (C-A: *AME* = 0.10, *p* = 0.078). By the end of training, bees performed above chance for all three stimulus types (Figure 4 C; Figure S2 B; A: mean = 0.63, CI = [0.52, 0.73], *p* = 0.008; B: mean = 0.61, CI = [0.49, 0.72], *p* = 0.024; C: mean = 0.72, CI = [0.61, 0.81], *p* < 0.001).

**Fig 4.**
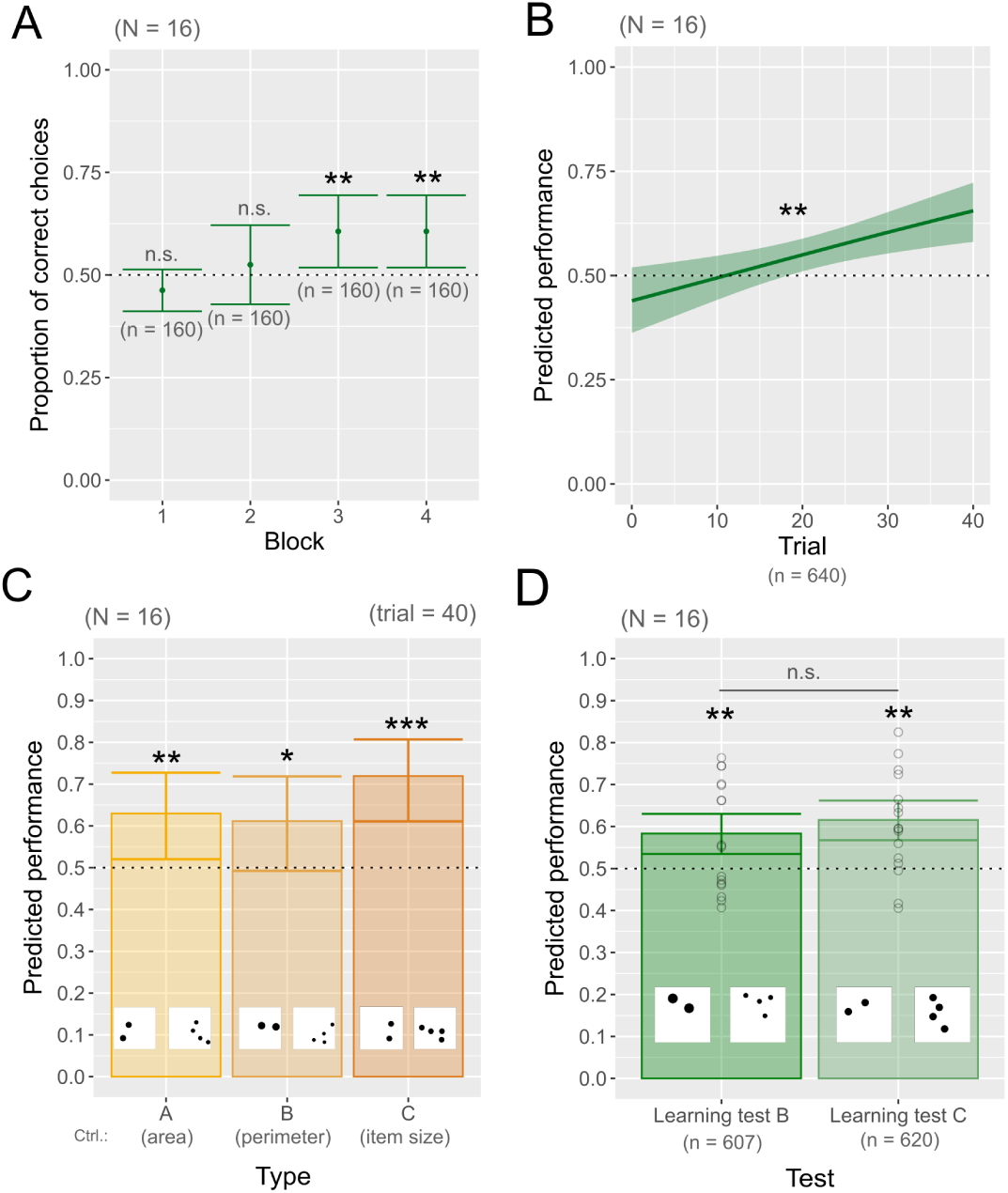
Experiment Space: Bees learned the quantity discrimination task. A) Mean proportion of correct choices across training blocks (Number of individuals: N = 16; Number of observations: n = 640). Each point represents the mean performance for a block of 10 trials. Error bars denote 95% CIs. Values are computed from raw data. B) Predicted probability of making a correct choice across trials (N = 16, n = 640). The solid line represents the mean probability to make a correct choice at each trial. Shaded areas indicate 95% CIs. Data and significance are computed from model 2.0. C) Predicted performance in the last training trial (trial 40). Each bar represents the predicted mean. Error bars denote 95% CIs. Data and significance are computed from model 2.0. D) Predicted performance in the two learning tests (N = 16; n = 1459). LT_B corresponds to the test in which total perimeter was controlled, and LT_C to the test in which item size was controlled. Each point represents the performance of a bee in a single test bout (45 s), computed from raw data; data and significance are computed from model 2.1. The dotted horizontal line shows chance level (0.5). (***): p ≤ 0.001, (**): p ≤ 0.01, (*): p ≤ 0.05, (■): p ≤ 0.1, (n.s.): p > 0.1. Ctrl.: control; N: number of individuals; n: number of observations.

In the two learning tests, which controlled either total perimeter (LT_B) or item size (LT_C), bees again performed above chance (Figure 4 D; LT_B: mean = 0.62, CI = [0.57, 0.66], *p* < 0.001; LT_C: mean = 0.58, CI = [0.53, 0.63], *p* < 0.001), with no difference between the tests (*p* = 0.260). Thus, bees learned the discrimination between quantities of items irrespective of non-numerical manipulations.

In both experiments, half of the bees were trained to choose the two-dotted stimulus to receive a reward (2:CS+) and the other half were trained to choose the four-dotted stimulus to receive a reward (4:CS+). A consistent difference emerged between these training groups: bees trained to select the two dots (2:CS+) tended to perform better during acquisition, although learning rates were comparable and all groups reached above-chance performance by the end of training. In Experiment Size, bees trained to select two dots significantly outperformed bees trained to select four dots (group[2-4]: *AME* = −0.09, *p* = 0.021; trial × group: *p* = 0.487; 2:CS+: mean = 0.75, CI = [0.67, 0.81]; 4:CS+: mean = 0.67, CI = [0.59, 0.74]). A similar pattern occurred in Experiment Space (Figure S3 A; Figure S4 A; group[2-4]: *AME* = −0.08, *p* = 0.050; trial × group: *p* = 0.131; 2:CS+: mean = 0.69, CI = [0.61, 0.76]; 4:CS+: mean = 0.62, CI = [0.53, 0.69]). During the learning test, both groups performed significantly above chance, but the direction of the group difference varied between experiments. In Experiment Size, bees trained to choose 4 dots showed a trend towards better performance (group[2-4]: *AME* = 0.06, *p* = 0.068; 2:CS+: mean = 0.57, CI = [0.52, 0.62]; 4:CS+: mean = 0.64, CI = [0.59, 0.68]). In Experiment Space, the opposite pattern emerged, with bees trained for two dots performing significantly better (Figure S3 B; Figure S4 B; group[2-4]: *AME* = −0.08, *p* = 0.047; 2:CS+: mean = 0.64, CI = [0.59, 0.69]; 4:CS+: mean = 0.56, CI = [0.50, 0.61]). Overall, these results indicate that the magnitude of the rewarded quantity influenced absolute performance levels but did not alter learning dynamics.

### Bees spontaneously encode numerousness

In the numerical transfer tests (TN), we homogenized the non-numerical cue that was predictive during training (total area in Experiment Size; convex hull in Experiment Space) so that it was no longer informative while numerosity (2 vs. 4) remained unchanged (Figure 5). Bees successfully solved this transfer test in both experiments (Figure 6 A,B; Experiment Size: mean = 0.61, CI = [0.58, 0.65], *p* ă 0.001; Experiment Space: mean = 0.60, CI = [0.55, 0.65], *p* ă 0.001). These results indicate that bees spontaneously encoded numerousness, even when non-numerical cues reliably predicted the reward during training.

**Fig 5.**
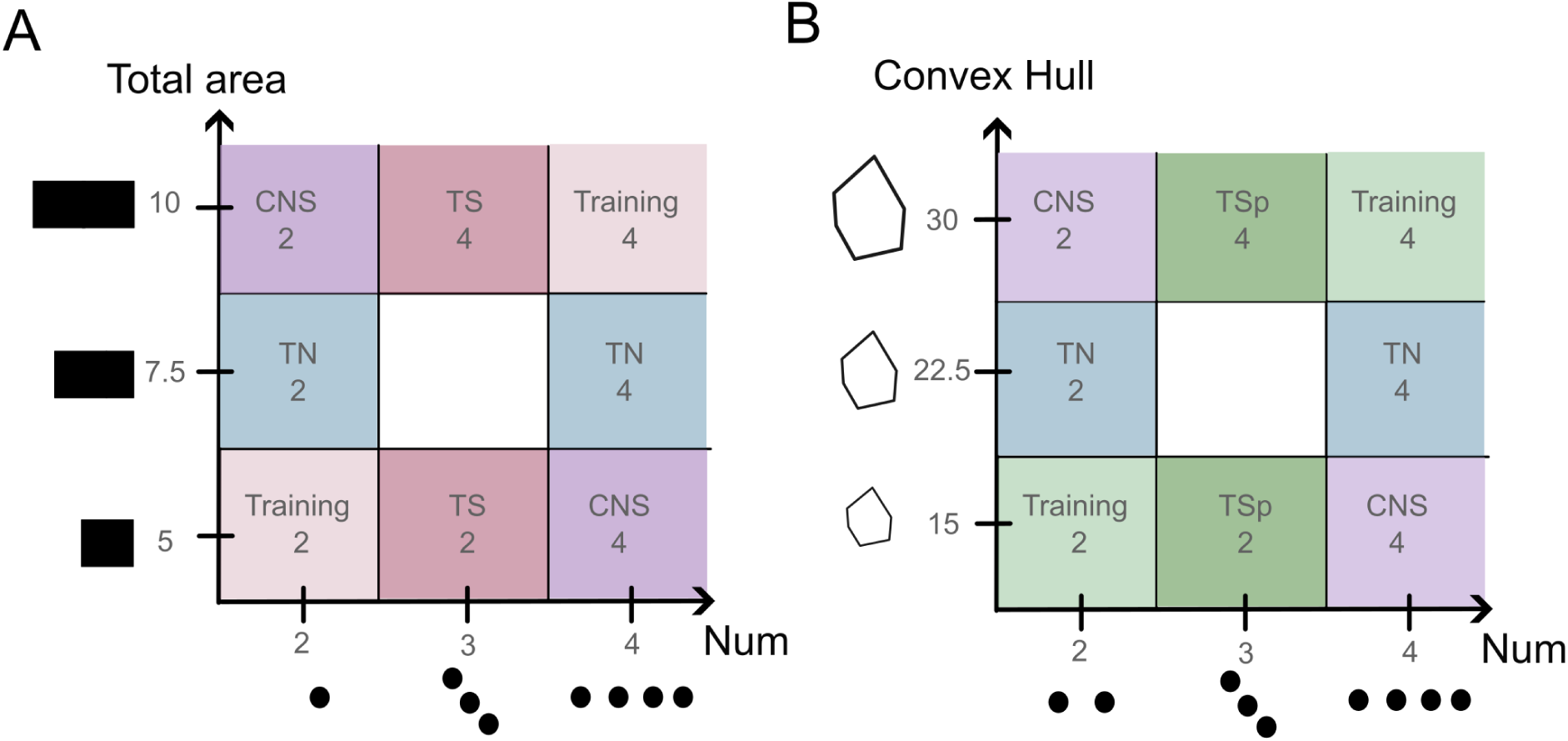
Matrix of training and testing stimuli overview. A) Representation of numerosity and total area values for the stimuli used in Experiment Size. On the x-axis are shown the three numerosity values used to design the stimuli in this experiment (2, 3, and 4); on the y-axis are shown the three total-area values (5, 7.5, and 10 cm^2^). Each cell represents a stimulus type defined by its numerosity and total area. Stimuli used during training correspond to the light-pink positions, while stimuli used in the numerical transfer test appear in blue, those used in the Size transfer test in dark pink, and those used in the conflict test in purple. B) Representation of the value of numerosity and convex hull area for the stimuli presented in the training (in light green), numerical transfer (in blue), Space transfer (in dark green) and conflict (in purple) tests in Experiment Space. CNS: conflict test, TN: numerical transfer, TS: Size transfer, TSp: Space transfer.

**Fig 6.**
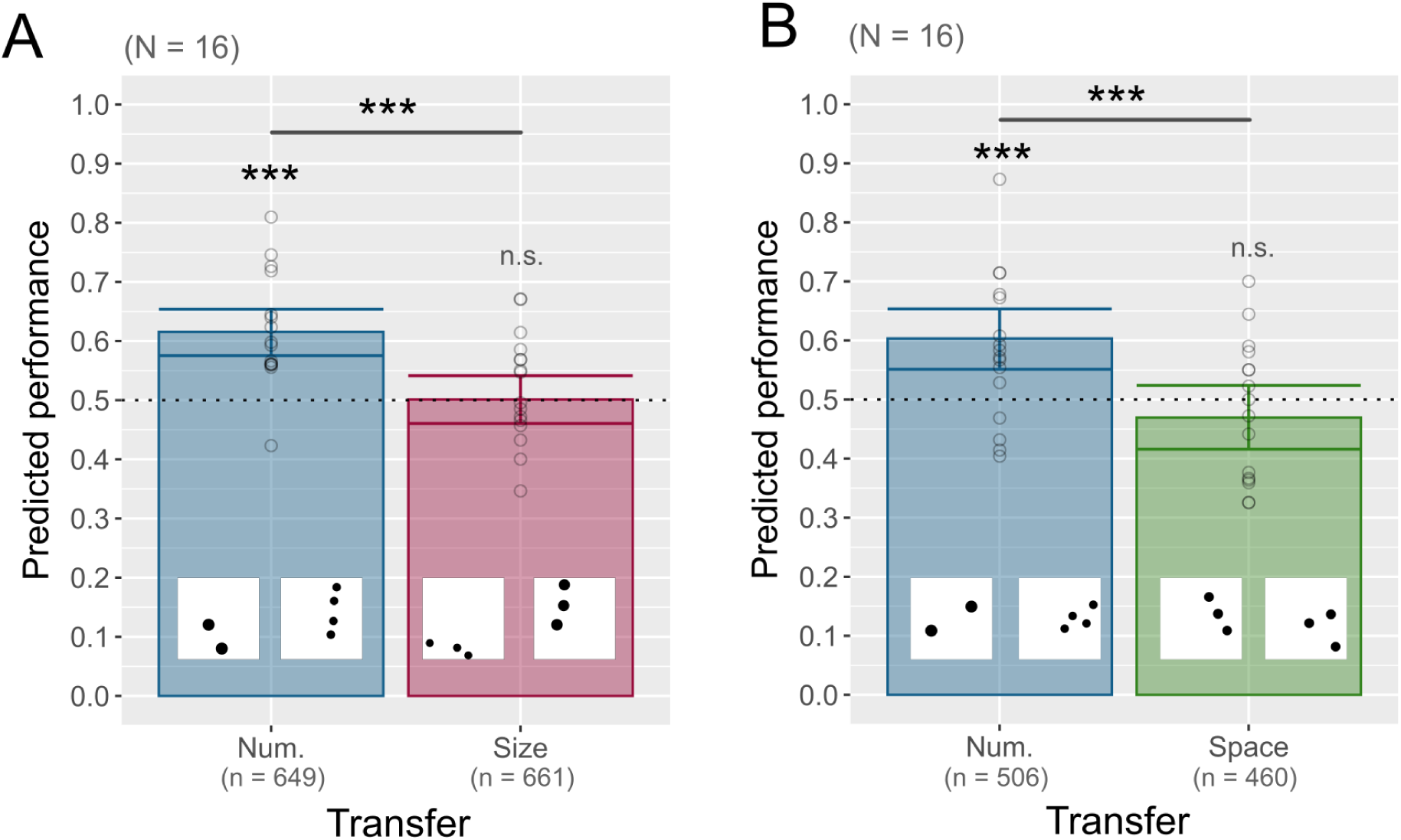
Bees spontaneously encode and favor numerousness. A) Predicted performance in the transfer tests of Experiment Size (N = 16, n = 1290). In the Numerical transfer (TN), total area was homogenized and therefore uninformative. In the Size transfer (TS), numerosity was homogenized. Predictions and significance are computed from model 1.2. B) Predicted performance in the transfer tests of Experiment Space (N = 16, n = 966). In the Numerical transfer (TN), convex hull was homogenized. In the Space transfer (TSp), numerosity was homogenized. Each bar shows the predicted mean. Error bars denote 95% CIs. Each point represents the performance of one bee in a single 45 s test bout (raw data). The dotted line indicates chance level (0.5). (***): p ≤ 0.001, (**): p ≤ 0.01, (*): p ≤ 0.05, (■): p ≤ 0.1, (n.s.): p > 0.1. N: number of individuals; n: number of observations.

**Fig 7.**
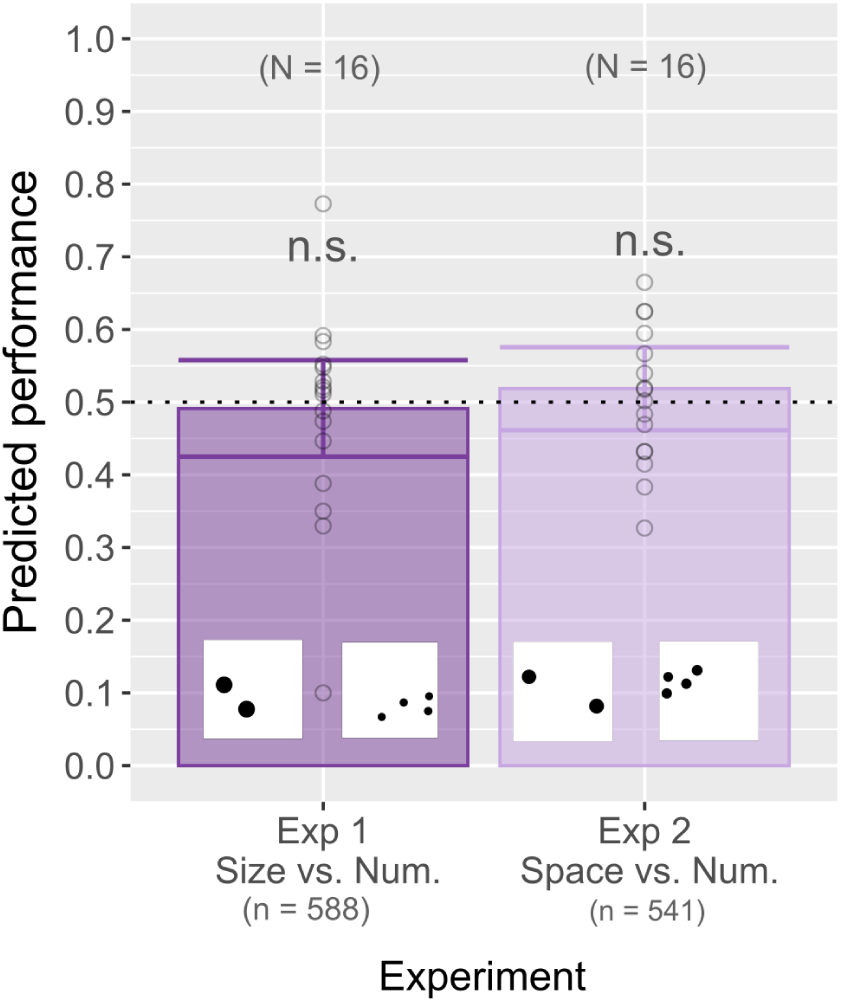
Bees did not show preference in the conflict test at the group level. A) Predicted performance in the conflict test of Experiment Size (N = 16, n = 566). B) Predicted performance in the conflict test of Experiment Space (N = 16, n = 541). Each bar shows the predicted mean, and error bars denote 95% CIs. Each point represents the mean performance of one bee in a single test bout (raw data). Data and significance for Experiment Size come from model 1.3; data and significance for Experiment Space come from model 2.4. The dotted horizontal line indicates chance level (0.5). (***): p ≤ 0.001, (**): p ≤ 0.01, (*): p ≤ 0.05, (■): p ≤ 0.1, (n.s.): p > 0.1. N: number of individuals; n: number of observations.

### Bees favor numerousness

In the non-numerical transfer tests (Size Transfer in Experiment Size; Space Transfer in Experiment Space), stimuli differed only in the relevant non-numerical cue (total area in Experiment Size; convex hull in Experiment Space), while numerosity was homogenized. Under these conditions, bees performed at chance level (Figure 6 A,B; Experiment Size: mean = 0.50, CI = [0.46, 0.54], *p* = 0.960; Experiment Space: mean = 0.47, CI = [0.42, 0.52], *p* = 0.274), and their performance was significantly lower than in the numerical transfer tests (Experiment Size: *p* ă 0.001; Experiment Space: *p* ă 0.001). These results indicate that bees relied primarily on numerosity during training rather than on non-numerical cues.

### Bees’ strategies: a numerical bias and a generalist strategy

In the conflict test (CNS), numerical and non-numerical cues were both available but pitted against each other. A performance above chance level indicates that bees rather choose the “numerically correct” stimulus, while a performance below chance level corresponds to choices made toward the “non-numerical correct” stimulus.

At the group level, bees performed at chance level (Exp. Size: mean = 0.49, CI = [0.42, 0.56], *p* = 0.798; Exp. Space: mean = 0.52, CI = [0.46, 0.57], *p* = 0.522; Figure 8).

**Fig 8.**
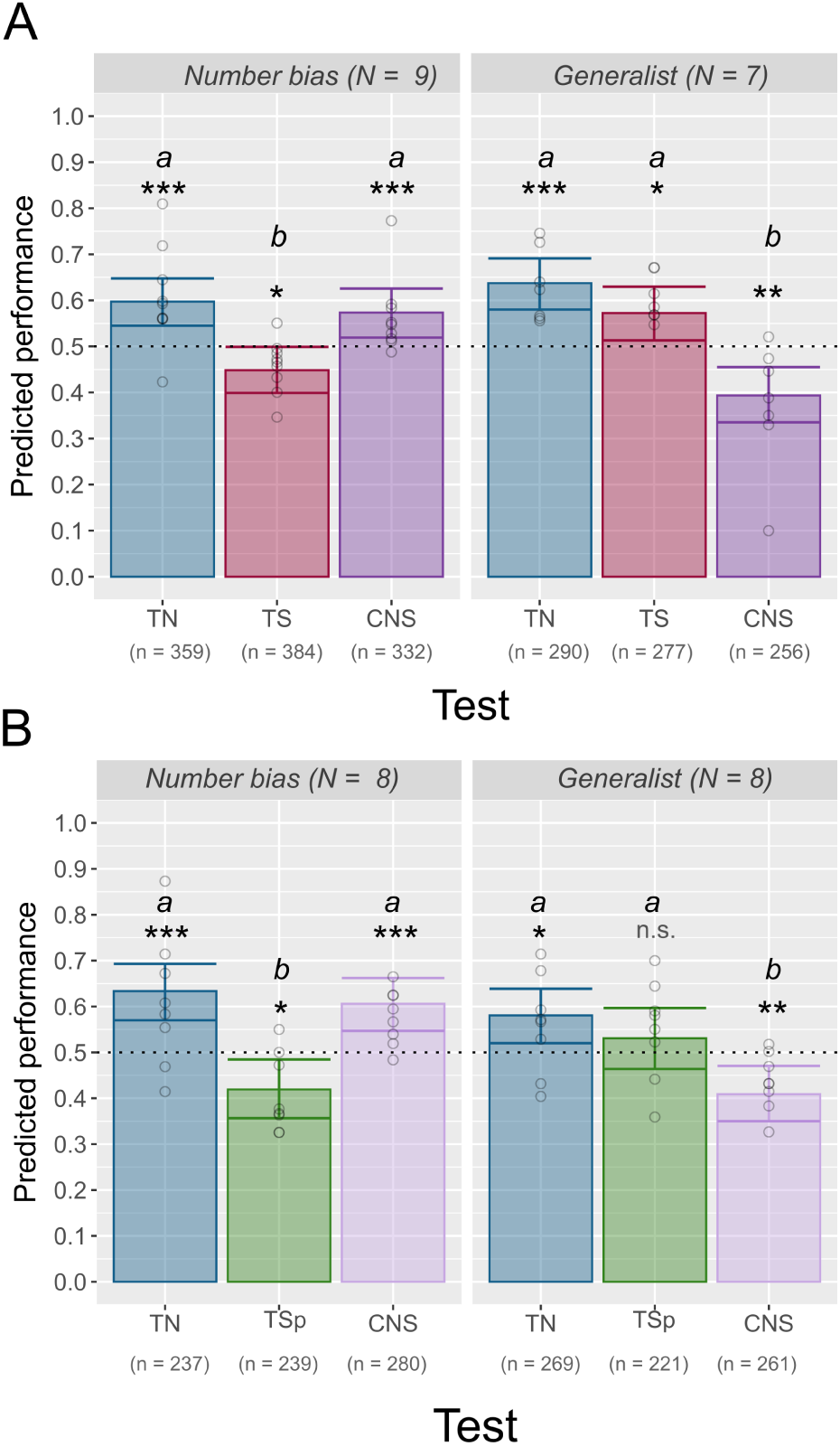
Bees’ strategies: a number bias and a generalist strategy. Experiment Size: A) Predicted performance in transfer and conflict tests for bees in the first cluster using a “number bias” strategy (N = 9, n = 1075). B) Predicted performance in transfer and conflict tests for bees in the second cluster using a “generalist” strategy (N = 7, n = 823). Data and significance for Experiment Size come from model 1.4. Experiment Space: C) Predicted performance in transfer and conflict tests for bees in the first cluster using a “number bias” strategy (N = 8, n = 756). D) Predicted performance in transfer and conflict tests for bees in the second cluster using a “generalist” strategy (N = 8, n = 751). Data and significance for Experiment Space come from model 2.5. Each bar shows the predicted mean, and error bars denote 95% CIs. Each point represents the mean performance of one bee in a single test bout (raw data). The dotted horizontal line indicates chance level (0.5). Different letters indicate significant pairwise differences between groups (holm-adjusted, p < 0.05). (***): p ≤ 0.001, (**): p ≤ 0.01, (*): p ≤ 0.05, (■): p ≤ 0.1, (n.s.): p > 0.1. CNS: conflict test; N: number of individuals; n: number of observations; Num.: numerosity; TN: numerical transfer; TS: Size transfer; TSp: Space transfer.

However, our cluster analysis revealed that this seemingly null effect masked two subgroups of bees, present in both experiments, that adopted opposite decision strategies.

In Experiment Size, the first cluster (N = 9; Figure 8 A) showed a clear numerical bias strategy. Bees’ performance remained similar whenever numerosity was available, in both the numerical transfer and conflict tests, and dropped only when numerosity was removed in the Size transfer (TN-CNS: *p* = 0.576; TS-TN: *p* ă 0.001; TS-CNS: *p* = 0.002). The bees in this subgroup succeeded in the numerical transfer, dropped below chance in the Size transfer, and significantly preferred the stimulus presenting the correct numerosity in the conflict test (TN: mean = 0.58, CI = [0.52, 0.65], *p* = 0.003; TS: mean = 0.44, CI = [0.38, 0.50], *p* = 0.0316; CNS: mean = 0.57, CI = [0.50, 0.63], *p* = 0.0316). The second cluster (N = 7; Figure 8 B) followed a more generalist strategy. Performance in both the numerical transfer and Size transfer was above chance and did not significantly differ (TN: mean = 0.64, CI = [0.57, 0.71], *p* ă 0.001; TS: mean = 0.58, CI = [0.51, 0.64], *p* = 0.010; TS-TN: *p* = 0.103). In the conflict test, performance dropped below chance, consistent with bees choosing according to total area (CNS: mean = 0.40, CI = [0.33, 0.48], *p* = 0.004; CNS-TN: *p* ă 0.001; CNS-TS: *p* ă 0.001). Experiment Space revealed a similar pattern. The first cluster (N = 8, Figure 8 C) again showed a numerical bias strategy, with comparable performance across the numerical transfer and conflict tests (TN-CNS: *p* = 0.566; TSp-TN: *p* ă 0.001; TSp-CNS: *p* ă 0.001), while performance in the Space transfer (TSp) dropped significantly below chance (TN: mean = 0.63, CI = [0.55, 0.70], *p* ă 0.001; TSp: mean = 0.42, CI = [0.34, 0.49], *p* = 0.015; CNS: mean = 0.61, CI = [0.53, 0.74], *p* ă 0.001). As in Experiment Size, these bees performed well whenever numerosity was available but not when only the non-numerical cue was informative. The second cluster (N = 8; Figure 8 D) resembled the generalist strategy, with similar performance across the numerical and Space transfers (TSp-TN: *p* = 0.266) and a drop toward the non-numerical cue in the conflict test (CNS-TN: *p* ă 0.001; CNS-TSp: *p* = 0.016). Here, the numerical transfer exceeded chance level (TN: mean = 0.58, CI = [0.51, 0.65], *p* = 0.018), the conflict test performance of this subgroup was below chance level (CNS: mean = 0.41, CI = [0.34, 0.48], *p* = 0.010), whereas the Space transfer was not significantly above chance level (TSp: mean = 0.53, CI = [0.45, 0.61], *p* = 0.377). Performance during the learning tests did not differ significantly between the two clusters in Experiments 1 and 2 (Figure S5 A,B; Figure S6; Experiment Size: *p* = 0.144; Experiment Space: *p* = 0.122). Together, these two clusters reveal inter-individual variability in *how*bees evaluate quantity, and which cue they rely on according to context.

### Mental number line congruence effect

Bees showed a left-to-right congruence effect: performance increased when the smaller quantity appeared on the left and the larger quantity on the right. This pattern was reflected in a significant side × group interaction, indicating that bees trained to “choose 2” (2:CS+) performed better when the correct stimulus was positioned on the left, whereas bees trained to “choose 4” (4:CS+) performed better when it appeared on the right. In Experiment Size, analyzing all test phases together (learning, transfer, and conflict tests) revealed a significant side × group interaction (Figure 9; *p* = 0.004). Bees in the 2:CS+ group performed significantly better when the correct stimulus was located on the left (side[R-L]: *AME* = 0.06, *p* = 0.016), whereas bees in the 4:CS+ group tended to perform better when it appeared on the right (side[R-L]: *AME* = −0.04, *p* = 0.088). This lateralization effect did not differ significantly across tests (side × group^ test: *p* = 0.103), and no such interaction was detected during training (side × group: *p* = 0.306). In Experiment Space, the overall testing analysis did not reveal a significant side × group interaction (*p* = 0.184). However, we found a significant interaction with test type (Figure S7; *p* = 0.003), indicating that the bias appeared selectively depending on the test. Bees trained to select two dots showed a significant left-to-right advantage in the Space transfer (*p* = 0.002), and bees in the 4:CS+ group showed a mirrored significant right-to-left advantage in the conflict test (*p* = 0.008). Together, these findings are consistent with a left-to-right spatial bias in quantity evaluation, particularly robust when bees must learn to differentiate quantities that covary in Size and Numerosity.

**Fig 9.**
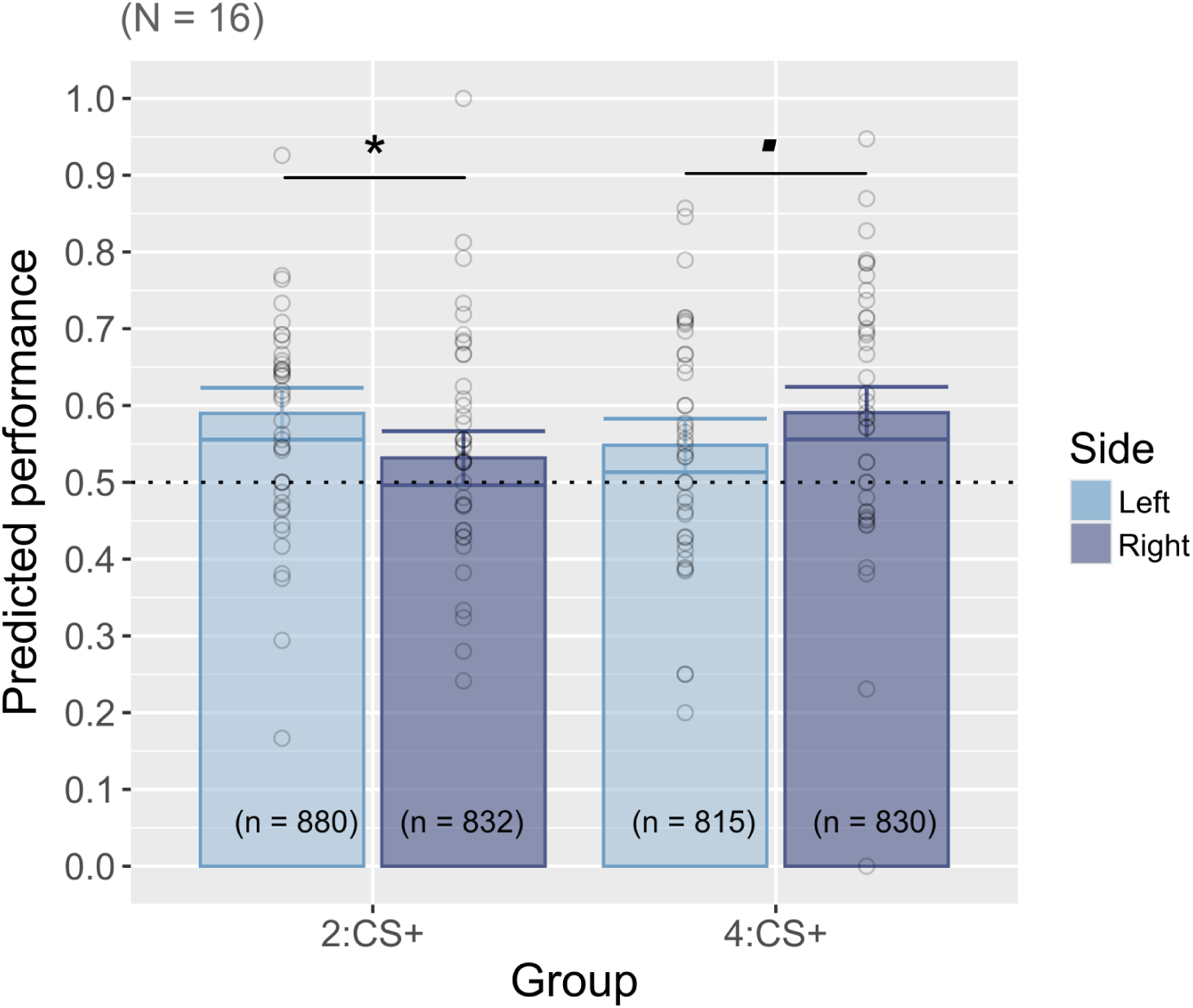
Mental number line congruence effect. Predicted performance in the testing phase of bees from Experiment Size for each group depending on the side of the correct stimulus. The 2:CS+ group is trained to “go toward 2”/“go to smaller” and the 4:CS+ group is trained to “go toward 4”/“go to larger”. Data and significance come from model 1.4. Each bar shows the predicted mean, and error bars denote 95% CIs. Each point represents the mean performance of one bee in a single test bout (raw data). The dotted horizontal line indicates chance level (0.5). (*): p ≤ 0.05, (■): p ≤ 0.1. N: number of individuals; n: number of observations.

### From spatial heuristics to rule learning

During training, bees faced a simple spatial choice: fly to the left or to the right arm of the Y-maze. An often overlooked feature of such paradigms is that, before extracting rules from the visual stimuli, bees may first rely on simple spatial heuristics, for example, returning to the side where they previously found a reward, regardless of the stimuli presented. We found that three key variables influenced bees’ choices during the training phase of Experiment Size: the arm in which the reward was previously obtained (Reward(*t*–1)), the arm chosen on the preceding trial (Choice_side(*t*–1)), and the side on which the correct stimulus appeared in the current trial (Correctp*t*q). All three variables significantly influenced bees’ choices differently across the training phase (Reward(*t*–1) × *block* : *p* = 0.005; Correctp*t*q × *block* : *p* = 0.030; Choice_side(*t*–1) × *block* : *p* = 0.069; Figure 10). In the first block (trials 2-10), bees significantly chose the same side as in the previous trial, reflecting a strong side bias (Choice_side(*t*–1) : *AME* = 0.26*, p* = 0.002). They also showed a tendency to return to the side where they had just obtained a reward in the previous trial (Reward(*t*–1) : *AME* = 0.15*, p* = 0.084). In the second block, bees still tended to favor the previously rewarded side (Reward(*t*–1) : *AME* = 0.15*, p* = 0.059), but the side bias disappeared (Choice_side(*t*–1) : *p* = 0.772). A strategy shift occurred in the third block: bees now significantly chose the side that was not rewarded in the last trial (Reward(*t*–1) : *AME* = 0.22*, p* = 0.010), and started to show a tendency to be influenced by the side where the correct stimulus was located (Correctp*t*q : *AME* = 0.15*, p* = 0.078). Only in the final block did bees consistently ignore spatial heuristics and rely on the stimuli themselves. The side of the correct stimulus had a strong effect on choice (Correctp*t*q : *AME* = 0.32*, p* ă 0.001), while both past reward and past side choice no longer influenced behavior (Reward(*t*-1) : *p* = 0.844; Choice_side(*t*-1) : *p* = 0.950).

**Fig 10.**
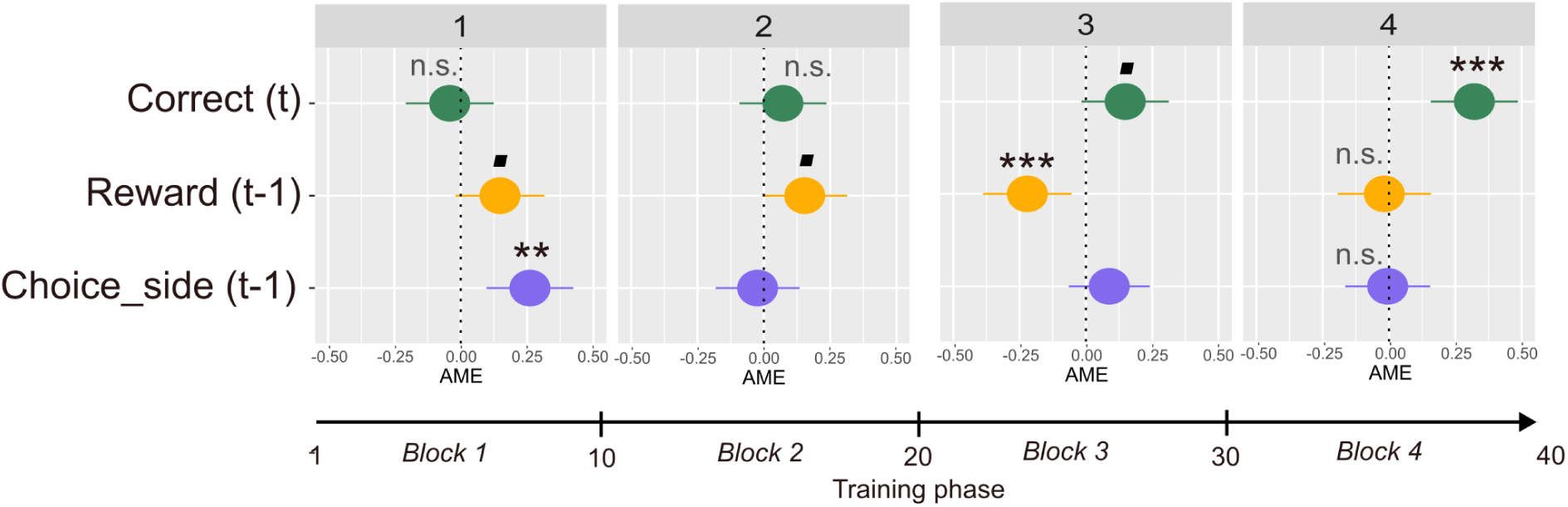
From heuristics to rule learning: navigating a Y-maze. Average marginal effects of three predictors on the side chosen by bees across the four training blocks. Reward(t–1) indicates the side where the reward was obtained on the previous trial; Choice_side(t–1) the side chosen on the previous trial; Correctptq the side where the correct stimulus was displayed. The vertical dashed line represents zero effect, corresponding to chance (no influence on side choice). (***): p ≤ 0.001, (**): p ≤ 0.01, (*): p ≤ 0.05, (■): p ≤ 0.1, (n.s.): p > 0.1. AME: Average marginal effect.

In Experiment Space, we found evidence that all three variables significantly influenced bees’ choices across the full training phase (Fig. SS8). Overall, bees showed a significant effect of the previously chosen side, consistent with a side bias that may differ across individuals (*AME* = 0.09, *p* = 0.024). Bees also tended to avoid the side where the reward had been obtained on the preceding trial (*AME* = −0.08, *p* = 0.062). Finally, and in line with our general analysis of training performance, bees were significantly influenced by the side on which the correct stimulus appeared (*AME* = 0.11, *p* = 0.010).

Overall, these patterns show that bees start by relying on simple spatial heuristics but gradually shift toward using the visual rule required by the task.

## Discussion

Recent work suggests that insects may possess a form of “number sense” despite their miniature brains (Bengochea et al., 2023; Bortot et al., 2021; Gatto et al., 2022; Giurfa, 2019; Howard et al., 2018, 2019a, 2019c; Nieder, 2021; Skorupski et al., 2018). In these studies, stimulus properties are manipulated so that non-numerical cues become less informative than numerousness for solving the task. A central question that follows is whether insects would still rely on numerousness when less computationally demanding cues are available to judge quantity. Here we show that, after being trained in conditions where numerical and non-numerical cues covary, honeybees succeed in quantity discrimination when numerousness was accessible but fail when only non-numerical cues were available. Inter-individual analyses further revealed two stable strategies: a “numerical bias” strategy, in which bees rely exclusively on numerousness, and a “generalist” strategy, in which bees encode multiple cues and preferentially follow non-numerical information in conflicting situations. Together, these results demonstrate that honeybees spontaneously encode numerousness and exhibit a population-level numerical bias, underpinned by consistent inter-individual differences in how quantity cues are favored.

### Bees spontaneously encode numerousness: a critical analysis of the influence of non-numerical cues

The ecological value of numerousness has long been debated (Leibovich et al., 2017; Núñez, 2017). In natural settings, animals encounter multiple quantity cues, and numerosity is only one of several dimensions available when comparing sets. This observation inspired the early “last-resort” hypothesis in non-human animals (Davis & Memmott, 1982; Davis & Pérusse, 1988), which proposed that animals would rely on numerosity only when no other cue remained available. Our results tell a different story. Across two complementary experiments in which numerical and non-numerical cues covaried during training, bees nevertheless extracted numerousness. Because our numerical transfer tests could not eliminate all non-numerical variation, we now examine whether any of these cues could plausibly account for the observed numerical bias.

#### Size vs. Numerousnes

In Experiment Size, bees successfully discriminated two from four items when numerical and Size cues (e.g., total area, total perimeter) covaried perfectly in the learning tests. In the subsequent numerical transfer test (TN), they continued to choose the correct numerosity even though the Size ratios experienced during training were no longer present. Several non-numerical cues still varied with numerosity in this test (Table 2). Each test in our protocol was designed such that if a bee relied on a specific non-numerical cue in the numerical transfer test, we could infer it from other tests. In the numerical transfer test, total area (r = 1), density (r = 1.1), and mean inter-item distance (r = 1) were essentially matched across numerosities, whereas total perimeter (r = 1.4), convex hull (r = 1.8), item size (r = 0.7), and total power (r = 0.66) remained correlated with numerosity (r = 2).

In regard to Size cues, inthe numerical transfer test, total area was homogenized. It was therefore not physically possible to control simultaneously the other Size cues, total perimeter and item size. Item size was larger in the 2-dotted stimuli than in the 4-dotted stimuli during TN. As item size was controlled in all stimuli during the training phase it is unlikely that bias could emerge from the training phase. However, bees could have been attracted to the stimulus presenting the larger items (as reported in humans, Lindskog et al., 2021). If so, bees rewarded to choose the 2 dots during the training should have choosed “correctly” but bees that were trained to choose the 4 dots should have failed this test, choosing the 2-dotted stimulus. This was, however, not the case in our data, as both training groups performed at similar levels. Another influence from the item size could have been a transfer from the “go to larger/go to smaller” total area during the training to “go to smaller/go to larger” item size during the numerical transfer (Bortot et al., 2020). This strategy would have led to bees choosing the “wrong” stimulus in each training group, bees trained to go toward 2 dots could have chosen the 4 dots and conversely. This was not the case.

On the other hand, during the numerical transfer test, the total perimeter of the 2 dotted stimulus was inferior to the one of the 4 dotted stimulus. This was also the case during the training phase. Bees possess strong edge-detection abilities (Kern et al., 1997; Lehrer & Srinivasan, 1993; Lehrer et al., 1990), the total perimeter is then a plausible candidate for guiding discrimination. To test this hypothesis, we used the same perimeter ratio in both the numerical transfer and the Size transfer test. If bees relied on total perimeter in the numerical transfer, they should have succeeded in both numerical and Size transfer. Instead, bees performed at chance in the Size transfer test at the population level. However, if we look at the cluster analysis, the “generalist” cluster (N = 7) succeeded in both of these transfer tests. Could the “generalist cluster” have used the total perimeter instead of numerousness all along? In the additional conflict test, the total perimeter was homogenized. Despite this inability to use the total perimeter, bees from the “generalist” cluster preferred the stimulus with the correct total area. Additionally, their performance did not track the decreasing predictive value of perimeter from learning to transfer tests. Under a non-numerical interpretation of these data, “generalist” bees would have to rely first on total area, then use total perimeter but only when area is uninformative, and additionally this hypothesis requires that bees would have a higher perceptual acuity for variation of perimeters than area. We thus consider this possibility as probably less parsimonious than considering that bees used numerousness as a cue.

Spacing cues (convex hull, density, and inter-distance) were homogenized over one-third of the training phase, however, the convex hull and the inter-distance remained predictable regarding their mean ratio. Importantly, total power was congruent with numerosity and Size cues over the whole training phase (Table 2). In the numerical transfer test, density (r = 1.1), and mean inter-item distance (r = 1), were non-predictive, spatial frequency total power was incongruent with numerosity (r = 0.66), but convex hull remained congruent with numerosity and could potentially have driven the bee’s choices (r = 1.8). Two lines of evidence argue against convex hull as a guiding cue. First, in Experiment Space, where convex hull covaried with numerosity in 100% of stimuli during training, bees chose randomly in the Space transfer test (TSp), despite the same predictive convex-hull ratio (r = 2). Second, in Experiment Size, bees performed better when inter-item distance rather than convex hull predicted numerosity, even though the convex-hull ratio was maximal in this condition (r = 2.8). Additionally, Shilat et al. (Shilat et al., 2021) recently emphasized that convex-hull shape should also be considered. In the numerical transfer test, both numerosities in the second stimulus set shared the same linear arrangement, and our analysis did not detect an effect of such equal convex-hull shape on performance. Additionally, total power (our spatial-frequency proxy) was incongruent with numerosity (training: r = 1.8; TN: r = 0.66), ruling it out as an explanatory cue. All together, even though we cannot exclude all possible non-numerical cues, the convergence of evidence strongly supports a single parsimonious interpretation: bees relied on numerosity during training and made a choice according to perceived numerousness in the numerical transfer test.

Our results differ from those reported for stingless bees (Eckert et al., 2022), likely due to several methodological differences between the studies, although species-specific capacities remain a possibility. In our protocol, stimuli were presented vertically, and each bee made its decision from a controlled, equidistant viewing point. In contrast, Eckert et al. displayed stimuli horizontally, trained and tested bees collectively, and allowed individuals to approach from multiple directions and distances. Under such conditions, the perceived Size of the stimuli at the moment of choice cannot be reliably estimated (Wehner, 1967). Moreover, their paradigm relied on an absolute appetitive discrimination in which Space cues were not controlled, further limiting comparability with our approach, which controlled for Space cues. Nonetheless, using harmonized protocols to directly contrast these species in future work may offer valuable insights into the evolutionary forces shaping numerical cognition in insects.

#### Space vs. Numerousness

In Experiment Space (Table 3), convex hull covaried with numerosity during training, yet bees again selected the correct numerosity in the numerical transfer test, even though the convex-hull ratio experienced during training was no longer available. As in Experiment Size, several non-numerical cues varied with numerosity in the numerical transfer test: total perimeter (r = 1.4), item size (r = 0.7), density (r = 2), inter-item distance (r = 0.49), spatial frequency total power (r = 0.7).

**Table 3.** Stimuli characteristics used during the training and testing phase of Experiment Space. Value of Numerosity, Size and Space cues of quantity in training and testing stimuli. In **bold**, are values covarying during the training phase. Ratios are expressed as the division of the value of the stimulus with the larger numerosity (or CH during TSp) by the value of the stimulus with the smaller numerosity. Each value is the mean value of all similar stimuli. CH: convex hull, CNS: conflict test, D: density, ID: mean inter-distance, IT: item size, LT_B: learning test category B, LT_C: learning test category C, Num.: numerosity, Power: power spectrum derived from Fourier transform without the DC component (freq = 0), TA: total surface, TN: Numerical transfer, TP: total perimeter, TSp: Space transfer.

| Exp. Space | Type | Num. |  | IT |  | TA |  | TP |  | D |  | CH |  | ID |  | Power |  |
| --- | --- | --- | --- | --- | --- | --- | --- | --- | --- | --- | --- | --- | --- | --- | --- | --- | --- |
|  |  | pair | ratio | pair | ratio | pair | ratio | pair | ratio | pair | ratio | pair | ratio | pair | ratio | pair | ratio |
| Training | A | 2 | 2 | 1.1 | 0.7 | 7.5 | 1 | 13.7 | 1.4 | 0.13 | 1 | 15 | 2 | 5.1 | 1 | 95.25 | 0.8 |
|  |  | 4 |  | 0.8 |  | 7.5 |  | 19.4 |  | 0.13 |  | 30.2 |  | 5.1 |  | 71.51 |  |
|  | B | 2 | 2 | 1.3 | 0.5 | 10.9 | 0.5 | 16.6 | 1 | 0.11 | 1.27 | 19.5 | 1.48 | 5.1 | 1 | 117.74 | 0.4 |
|  |  | 4 |  | 0.7 |  | 5.5 |  | 16.6 |  | 0.14 |  | 28.1 |  | 5.1 |  | 47.77 |  |
|  | C | 2 | 2 | 1 | 1 | 6.3 | 2 | 12.6 | 2 | 0.15 | 0.71 | 15 | 2.80 | 5.1 | 1 | 64.68 | 1.90 |
|  |  | 4 |  | 1 |  | 12.5 |  | 25.1 |  | 0.11 |  | 37.6 |  | 5.1 |  | 119.89 |  |
|  | Mean | 2 | 2 | 1.1 | 0.7 | 8.2 | 1.1 | 14.2 | 1.4 | 0.13 | 1.0 | 16.3 | 2.10 | 5.1 | 1 | 92.55 | 1.0 |
|  |  | 4 |  | 0.8 |  | 8.5 |  | 20.3 |  | 0.13 |  | 32.0 |  | 5.1 |  | 79.72 |  |
| Testing | LT_B | 2 | 2 | 1.3 | 0.5 | 10.9 | 0.5 | 16.6 | 1 | 0.11 | 1.21 | 19.0 | 1.6 | 5.1 | 1 | 130.91 | 0.3 |
|  |  | 4 |  | 0.7 |  | 5.5 |  | 16.6 |  | 0.12 |  | 31.5 |  | 5.1 |  | 43.78 |  |
|  | LT_C | 2 | 2 | 1 | 1 | 6.3 | 2 | 12.6 | 2 | 0.15 | 0.8 | 13.4 | 2.5 | 5.1 | 1 | 70.47 | 1.8 |
|  |  | 4 |  | 1 |  | 12.6 |  | 25.1 |  | 0.12 |  | 33.5 |  | 5.1 |  | 122.14 |  |
|  | TN | 2 | 2 | 1.9 | 0.7 | 7.5 | 1 | 13.7 | 1.4 | 0.09 | 2 | 22.5 | 1 | 8.6 | 0.49 | 98.05 | 0.7 |
|  |  | 4 |  | 0.8 |  | 7.5 |  | 19.4 |  | 1.18 |  | 22.5 |  | 4.2 |  | 71.54 |  |
|  | TSp | 3 | 1 | 0.9 | 1 | 7.5 | 1 | 16.8 | 1 | 0.20 | 0.5 | 15 | 2 | 4.3 | 1.79 | 74.54 | 1.1 |
|  |  | 3 |  | 0.9 |  | 7.5 |  | 16.8 |  | 0.10 |  | 30 |  | 7.7 |  | 80.40 |  |
|  | CNS | 2 | 2 | 1.9 | 0.7 | 7.5 | 1 | 13.7 | 1.4 | 0.07 | 2.95 | 28 | 0.6 | 11.1 | 0.33 | 91.38 | 0.8 |
|  |  | 4 |  | 0.8 |  | 7.5 |  | 19.4 |  | 0.21 |  | 19 |  | 3.6 |  | 69.59 |  |

Because the convex hull was identical across numerosities in the numerical transfer test, density necessarily doubled in the four-dot arrays. However, density was not a reliable predictor during training (mean r = 1), and if bees had relied on it in the numerical transfer test, they should have performed at or below chance level in Learning Test C (r = 0.8).

Instead, bees of both clusters performed above chance level. Inter-item distance is similarly implausible: it was strictly matched during training in each pair and therefore is unlikely to guide behavior consistently in the numerical transfer test. Spatial frequency total power varied across the whole training and was on average uninformative (r = 1), and would have predicted differential performance in both learning tests, which in this experiment was not the case.

The total perimeter ratio in the numerical transfer test (r = 1.4) matched that of Experiment Size’s transfer tests. We already discussed that even when the total perimeter is fully predictable in the training phase, bees seem to be unable to refer to such a total perimeter ratio to guide their choice. In this experiment, the total perimeter was additionally controlled during the training phase, making it unlikely to have drive behavior in this test and not in the Experiment Size. Item size can also be excluded based on learning-test performance, where no cue-consistent performance emerged.

As in Experiment Size, none of the available non-numerical cues provides a coherent explanation for bees’ performance. We therefore consider that the most parsimonious interpretation across both experiments is that honeybees based their decisions on numerosity.

#### Effect of the magnitude of the rewarded stimuli

In both experiments, we observed a bias toward choosing the two-dot stimulus during the training phase. This bias disappeared during testing in Experiment Size, but persisted in Experiment Space tests. Importantly, it did not prevent bees from learning the discrimination. A similar asymmetry was reported by Gross et al. (Gross et al., 2009), where honeybees performed better when matching two items than when matching three in a delayed match-to-sample task but not in other numerical discrimination tasks in honeybees (Bortot et al., 2019). Interestingly, human infants show the opposite pattern, exhibiting greater difficulty when discriminating smaller quantities (Estes, 1976). Chicks are also known for spontaneously preferring the larger set of objects (Rugani, Regolin, & Vallortigara, 2010).

### Bees spontaneously favor numerousness: a need for context in quantity bias studies

At the population level, bees dropped to chance level when presented with equal numerosities, even though the non-numerical cues used during training (e.g., total area or convex hull) were predictive of the correct stimulus. This contrasts with reports in pigeons, which succeed in comparable Size-transfer tests (Diaz & Wasserman, 2023; Kubo, 2022). Instead, our results resemble those from rhesus macaques, whose categorization responses depended on numerosity rather than total area when both cues covaried during training (Ferrigno et al., 2017).

#### Generalization decrement

In our study, the Size and Space transfer tests presented a novel numerosity of three items. This potentially introduced a generalization decrement (Guttman & Kalish, 1956). However, no such decrement appeared in the numerical transfer test, where equally novel total area and convex hull values were introduced. On the other hand, the dots were arranged either linearly or triangularly, without affecting the bees performance. This arrangement makes it unlikely that generalization decrement alone accounts for the decline in performance in the Size and Space transfer tests.

#### Perceptual and mathematical ratios: discriminability of numerical, Size, and Space cues

A key issue recently highlighted in “number bias” studies is that numerical and non-numerical cues are often equated using mathematical ratios rather than perceptual variability (Aulet & Lourenco, 2023). The finer the acuity for a given cue, the larger the

perceptual variability for the same mathematical ratio. When the acuity for numerosity and total area differs, as is the case in humans (Odic et al., 2013), an apparent numerical bias may simply arise because the perceptual change in numerosity is larger than that of area. Indeed, when Aulet and colleagues (Aulet & Lourenco, 2023) matched cues based on perceptual rather than mathematical variability, infants shifted toward a Size bias, whereas adults continued to rely on numerousness. Their study, however, did not test non-human primates, leaving open whether the numerical bias reported in macaques (Ferrigno et al., 2017) would persist under perceptually matched conditions.

In honeybees, available evidence suggests that acuity for area and numerousness is relatively similar, making mathematical and perceptual matching effectively equivalent. A recent study by Dyer et al. (in review) estimated a Weber fraction of approximately 0.24 for total area using an appetitive-aversive differential conditioning paradigm. For numerosity, recent work employing comparable procedures suggests a Weber fraction of around 0.20 (Howard et al., 2019b). With such closely aligned acuities, differences in perceptual variability are unlikely to account for the numerical bias observed in our experiments. That said, Weber fractions can vary substantially across protocols (Clayton et al., 2015; Kuzmina & Malykh, 2022; Price et al., 2012; Smets et al., 2015). Future work should therefore directly compare Size, Space, and numerical acuity under strictly matched training conditions (e.g., number of trials, apparatus, conditioning procedure) to confirm whether these magnitudes are indeed equivalent in acuity.

#### Congruency effect and the theory of magnitude

Quantity discrimination during both training and learning tests was sensitive to non-numerical cues, even when they did not remain fully congruent with the other cues across all trials of the training. In the learning test of Experiment Size, bees performed better when inter-item distance was predictive of the correct stimulus, even though this cue varied within the training set. In Experiment Space, performance during training was higher when total area and total perimeter were congruent with numerosity compared to conditions in which one of these cues was uninformative. In numerical cognition, such interactions are referred to as a congruency effect: discrimination improves when numerical and non-numerical cues are aligned and decreases when not (Szűcs et al., 2013). Congruency effects have been documented in humans (DeWind et al., 2015; Leibovich et al., 2017; Szűcs et al., 2013) and in several non-human species, including dolphins (Kilian et al., 2003), apes (Tomonaga, 2008; Vonk et al., 2014), fish (Agrillo et al., 2011), and birds (Emmerton, 1998). For instance, mosquitofish learn 2 vs. 3 discrimination more rapidly when Size and numerosity covary than when they must rely on a single cue (Agrillo et al., 2011). Most studies highlight the influence of Size cues, particularly total area. The congruency effect is central to the “theory of magnitude” (ATOM) (Walsh, 2003, 2014), which proposes that magnitudes such as time, space, and numerosity share a common representational system (Davis & Pérusse, 1988). Under this framework, interactions between magnitudes arise naturally from a single processing mechanism underlying different forms of magnitude evaluation. ATOM is often invoked to argue against the existence of a dedicated “number sense” (Leibovich et al., 2017). Here, we argue instead that numerosity constitutes one dimension of quantity evaluation: it can be influenced by non-numerical cues, but it is not necessarily confounded with them. Our findings align with those of Bortot et al., who reported spontaneous transfer between Size and numerousness in honeybees (Bortot & Vallortigara, 2023; Bortot et al., 2020).

#### Contextualizing number bias

Although our results reveal a general numerical bias in honeybees, it is important to recognize that quantity bias can depend on both perceptual and ecological contexts (Agrillo, 2015). Ecologically, animals may weigh different quantity dimensions according to their adaptive value in a given situation. For example, assigning a higher weight (*w*) to total area when evaluating food resources, but not when assessing groups of conspecifics. Perceptually, cues should also receive greater weight when they offer higher discriminability (Grasso et al., 2022), such as relying on density rather than numerosity when processing highly textured visual stimuli (Anobile et al., 2014; Pomè et al., 2019). Thus, the numerical bias reported here should be interpreted within the specific conditions of our experiments, where numerical and non-numerical cues were matched in ratio and ecological relevance.

### Inter-individual variability: both a pure numerical bias and a generalist strategy coexist in bees’ population

When studying the cognitive abilities of an animal species, population-level results often receive most of the attention, and this may also be the case for our study. However, experiments frequently report substantial inter-individual variability, not only in the magnitude of cognitive performance but also in the strategies that individuals use to solve a task.

#### Early findings of bees inter-individual variability in their use of quantity cues

The first study to suggest proto-counting skills in honeybees already revealed clear inter-individual differences in how bees used quantity cues. In this experiment, Chittka and colleagues (Chittka & Geiger, 1995) trained foragers to visit a gravity-feeder providing sucrose solution after flying past three yellow tents placed along the route from the hive. Most bees solved the task by relying on the total distance traveled, using it as their primary cue to anticipate the reward location. However, about 20% of individuals behaved differently: they appeared to use the tents themselves as sequential landmarks. When the distance between tents was increased, these bees continued searching beyond the usual reward location, as if “enumerating” their passage through the first, second, and third tents, even though the overall distance no longer matched the trained value. Importantly, the proportion of bees using this apparent numerical strategy depended on the total discrepancy between the distance flown and the number of landmarks encountered. These early findings were later tested under more controlled conditions in which distance cues were rendered unusable. Under these constraints, the majority of bees were able to rely on the numerical order of landmarks when this was the only informative cue available (Dacke & Srinivasan, 2008).

#### A “numerical bias” strategy and a “generalist” strategy

Our data support the early evidence of Chittka and colleagues (Chittka & Geiger, 1995) that, when given the choice, bees may actually show inter-individual differences in cues they favor to resolve a cognitive task. The first strategy we described is a “numerical bias” strategy. In our data, this accounted for about 50 % of our tested bee population in each experiment. These bees succeeded whenever numerousness was available (numerical transfer and conflict tests) but dropped just below chance level when only non-numerical cues were available (Size and Space transfer). Their profile suggests that some individuals relied almost exclusively on numerousness, largely ignoring the non-numerical cues that covaried with numerousness during training, and were unaffected by incongruency when non-numerical cues and numerical ones were in discrepancy. However, we do not have a hypothesis explaining why those bees seemed to prefer the stimulus presenting the incorrect value of the non-numerical cues in the non-numerical transfer test. The second strategy followed a more “generalist” pattern. These bees appeared to succeed in both, the numerical and non-numerical transfer tests, indicating that they had learned from multiple dimensions during training. However, when the cues were placed in opposition, these individuals tended to follow the non-numerical cue.

While these strategies arise from exploratory analyses, their recurrence across both experiments makes the discovery difficult to dismiss and adds to the early-on stated need for an inter-individual view on insects’ numerical cognition (Pahl et al., 2013). More importantly, these findings highlight an often-overlooked reality: numerical cognition is rarely uniform within a species. Individuals can differ, sometimes profoundly, in how they weigh cues, extract rules, and solve quantitative problems. Whether these inter-individual differences are stable in time and context remains to be explored. If stable, these profiles could help resolve the long standing question of the function of numerical cognition across the animal realm, and especially in insects (i.e., “why”).

#### Individual differences in the use of quantity cues within other animal species

In other species, comparable inter-individual variability arises. In pigeons, some individuals solve numerical conflict tests by following numerosity cues, whereas others drop to chance level, which made the population-level score not significantly different from chance Kubo, 2022. In cats, two out of four subjects performed above chance in the numerical transfer test, yet the non-significant population result (p = 0.12) has often been taken as evidence that cats lack spontaneous numerousness encoding (Pisa & Agrillo, 2009). In both cases, population averages obscured meaningful individual strategies. Such variation was also reported in the use of non-numerical cues in fish, with some individuals favoring total area while others favored the cue of convex hull (Agrillo et al., 2009).

#### Potential mechanisms of inter-individual variability in the “numerical bias” in honeybees

In humans, inhibitory control is thought to underlie major inter-individual differences in numerical competence and following mathematical performance (Casanova et al., 2025; Szűcs et al., 2013). Numerousness becomes the preferred cue for evaluating quantity only once individuals learn to suppress competing signals from non-numerical dimensions. This inhibitory mechanism is considered central to the developmental shift observed between infants and adults, in which numerical information increasingly dominates behavioral choices under incongruent conditions (Rodríguez & Ferreira, 2023), precisely the type of situation that is recreated in our conflict tests.

In honeybees, workers are often treated as a homogeneous group, yet colonies contain individuals with distinct ecological specializations. Scouts and recruits, for instance, face different navigational demands and differ in their exploration-exploitation balance (Seeley, 1983). These behavioral roles are accompanied by cognitive asymmetries: scouts exhibit stronger latent inhibition than recruits, a non-associative process through which individuals learn to ignore irrelevant or previously encountered stimuli (Cook et al., 2019). Such variation offers a potential framework for interpreting the two strategies we identified, the “numerical-bias” and “generalist” clusters. If individuals differ in latent inhibition and/or in their ecological role within the colony, they may also differ in the extent to which they can suppress non-numerical cues or flexibly integrate multiple cues during quantity evaluation. Understanding how ecological specialization and cognitive traits map onto numerical strategies represents a promising direction for future work and may help clarify the ecological functions of numerousness processing in non-human animals.

### From left-to-right to few-to-many congruency effect

#### An innate and shared orientation of the SNA: ongoing debates

The first evidence for a left-to-right Spatial-Numerical Association (SNA) in humans arose unexpectedly: in parity judgments, participants responded faster to small numbers with the left hand and to large numbers with the right (Dehaene et al., 1993). This effect (SNARC), later replicated across tasks and modalities (Bulf et al., 2016; De Hevia et al., 2014; Fischer et al., 2003; Zorzi et al., 2002), suggests a systematic spatial mapping of numerical magnitude. Its origin, however, remains debated. One view attributes SNA orientation to cultural habits such as reading direction (Bulut et al., 2023; Shaki & Fischer, 2008; Zebian, 2005). Others report similar biases across populations with different writing systems, pointing instead to biological constraints (Eccher et al., 2025; Hochman et al., 2024; Pitt et al., 2021; Zohar-Shai et al., 2017). Findings in newborn infants and various non-human species further suggest that an oriented SNA may emerge independently of cultural experience (de Hevia et al., 2017; Giurfa, 2022; Rugani et al., 2015, 2020, 2024). Yet contradictory evidence persists: several primate, bird, and fish species show reversed or absent SNARC-like effects (Beran et al., 2019; Drucker & Brannon, 2014; Greenacre et al., 2022; Jackson et al., 2025; Rugani, Kelly, et al., 2010; Rugani et al., 2011; Triki & Bshary, 2018). Thus, whether a left-to-right SNA is shared and biologically rooted remains an open question—both in non-human animals (Beran et al., 2019; Jackson et al., 2025; Pitt et al., 2023; Rugani, Kelly, et al., 2010; Rugani et al., 2011; Triki & Bshary, 2018) and even in humans (Bulut et al., 2023; Eccher et al., 2025; Pitt et al., 2021, 2023; Shaki & Fischer, 2008).

#### SNA congruency effect in honeybees

Recent evidence suggests that honeybees exhibit a left-to-right spatial-numerical association (SNA), or Mental Number Line (MNL) (Giurfa et al., 2022; Kuo et al., 2025). Relative to a reference numerousness (e.g., 5), smaller numerosities (e.g., 3) shift honeybees’ attention toward the left visual field, whereas larger ones (e.g., 8) bias attention toward the right (Giurfa et al., 2022).

Beyond attentional shifts, humans and domestic chicks also show a left-right congruency effect: individuals perform better when small numerosities appear on the left and larger ones on the right (Loconsole et al., 2023). Our results reveal a comparable effect in honeybees. The bias was consistent in the testing phase of Experiment Size, but less stable across Experiment Space, suggesting a small, context-dependent influence on quantity discrimination, as suggested by recent evidence of the congruency effect in chicks (Loconsole et al., 2023). Taken together, our findings extend the left-to-right congruency effect beyond vertebrates and align with recent work indicating the presence of an SNA in honeybees (Giurfa et al., 2022).

#### Valence hypothesis of the oriented SNA?

Our study also provides insight into mechanistic hypotheses proposed to explain the biological origin of an oriented SNA, particularly the “valence hypothesis” (Vallortigara, 2018). This hypothesis links the SNA to emotional lateralization: smaller-than-expected numerosities would evoke negative valence (associated with right-hemisphere processing in humans (Davidson, 2004; Palomero-Gallagher & Amunts, 2022)), whereas larger numerosities would evoke positive valence (rather linked to the left hemisphere). Although grounded in human neurobiology, this framework is broadly consistent with the context-dependent nature of SNAs reported in other animals (Giurfa et al., 2022; Rugani et al., 2015).

However, the valence hypothesis does not align with our results. In our experiment, bees were differentially conditioned: some associated the smaller numerosity (2) with reward and the larger numerosity (4) with punishment, whereas others learned the opposite contingency. Under such training, it would be implausible for the presentation of “two dots” in the learning tests, for example, to evoke a negative-valence response in bees trained to get a reward at the two-dotted stimuli. The persistence of a left-to-right congruency effect across opposite reinforcement contingencies therefore argues against valence-based explanations, at least in honeybees.

Although investigating the origin of SNA orientation was not the primary goal of this study, our findings suggest that alternative hypotheses deserve further examination. One possibility is the “brain lateralization hypothesis”, which proposes that dominance of the right hemisphere in visual processing biases initial exploration toward the left visual field (De Hevia et al., 2014; Rugani et al., 2007; Vogel et al., 2003). Lateralization in honeybees remains poorly understood (Baracchi et al., 2018; Frasnelli et al., 2014; Rigosi et al., 2015). Future work on honeybees visual processing lateralization in non-numerical tasks could help orient the underlying mechanisms of the MNL. Additionally, future work targeting non-visual quantity evaluation may help distinguish if the SNA is linked to visuospatial machinery itself or is rather a property of quantity evaluation.

### Mechanistic perspective on rule learning in honeybees

The Y-maze is a classical tool in honeybee cognition research (Avargues-Weber et al., 2015; Avarguès-Weber, Portelli, et al., 2010; Avarguès-Weber et al., 2011, 2012; Bortot & Vallortigara, 2023; Bortot et al., 2019, 2020; Giurfa et al., 2001; Howard et al., 2019a, 2019b, 2019c), however, surprisingly little is known about the strategies bees deploy while learning within this apparatus. Building on this gap in knowledge, we analyzed the temporal sequence of honeybee choice during our learning tasks to test whether bees progressed through distinct learning phases. Our analysis implies the use of spatial heuristics on top of stimuli-guided learning in both experiments. In Experiment Size, we show a transition from location-based heuristics within the Y-maze to stimulus-based rule learning. Bees began by repeatedly choosing the same side, then switched to an alternation strategy in which they preferentially chose the side that was not rewarded at the previous trial, before finally showing consistent stimulus-based choices. This pattern hints at the possibility of hypothesis testing during learning tasks in honeybees (Wason, 1960). The predominance of a localization-based rule is consistent with honeybee foraging ecology. Under natural conditions, bees rely first on vector-based navigation, analogous to a simple “enter the maze and turn left” rule. Only when vector information becomes unreliable do bees seem to additionally use landmarks (Kheradmand & Nieh, 2019; Towne et al., 2017).

By capturing these transitions, our study provides a first glimpse into the temporal architecture of rule acquisition in honeybees and lays the groundwork for future computational models of conceptual learning in insects during Y-maze like dual force choices. A promising candidate is the use of bandit models within the reinforcement learning framework (Claeys et al., 2025). Reinforcement learning is already used as a tool to study insect navigation models (Lochner et al., 2024) and could help identify different sequential strategies and the potential for bees to switch between exploitation/exploration rates.

## Conclusion

Taken together, our findings show that honeybees spontaneously extract numerousness even when multiple quantity cues are available and appear to favor numerousness cues over Size and Space cues. We found that this numerical bias has emerged within an insect species possessing a brain of fewer than one million neurons, built on an architecture that bears little resemblance to the mammalian neocortex. Our results challenge the assumption that numerical representations require complex neural machinery. Instead of viewing insect numerical cognition as a simplified version of vertebrate systems, our findings support the idea that similar computational solutions for evaluating quantity may have evolved independently across distant lineages. Our findings and conclusions raise the possibility that numerical abilities shared across vertebrates could be analogous rather than homologous traits present in invertebrates as well. Future work exploring inter-individual variability and cross-species comparisons will be essential for uncovering the ecological problems for which numerousness may offer a selective advantage.

---

**Table S1.**
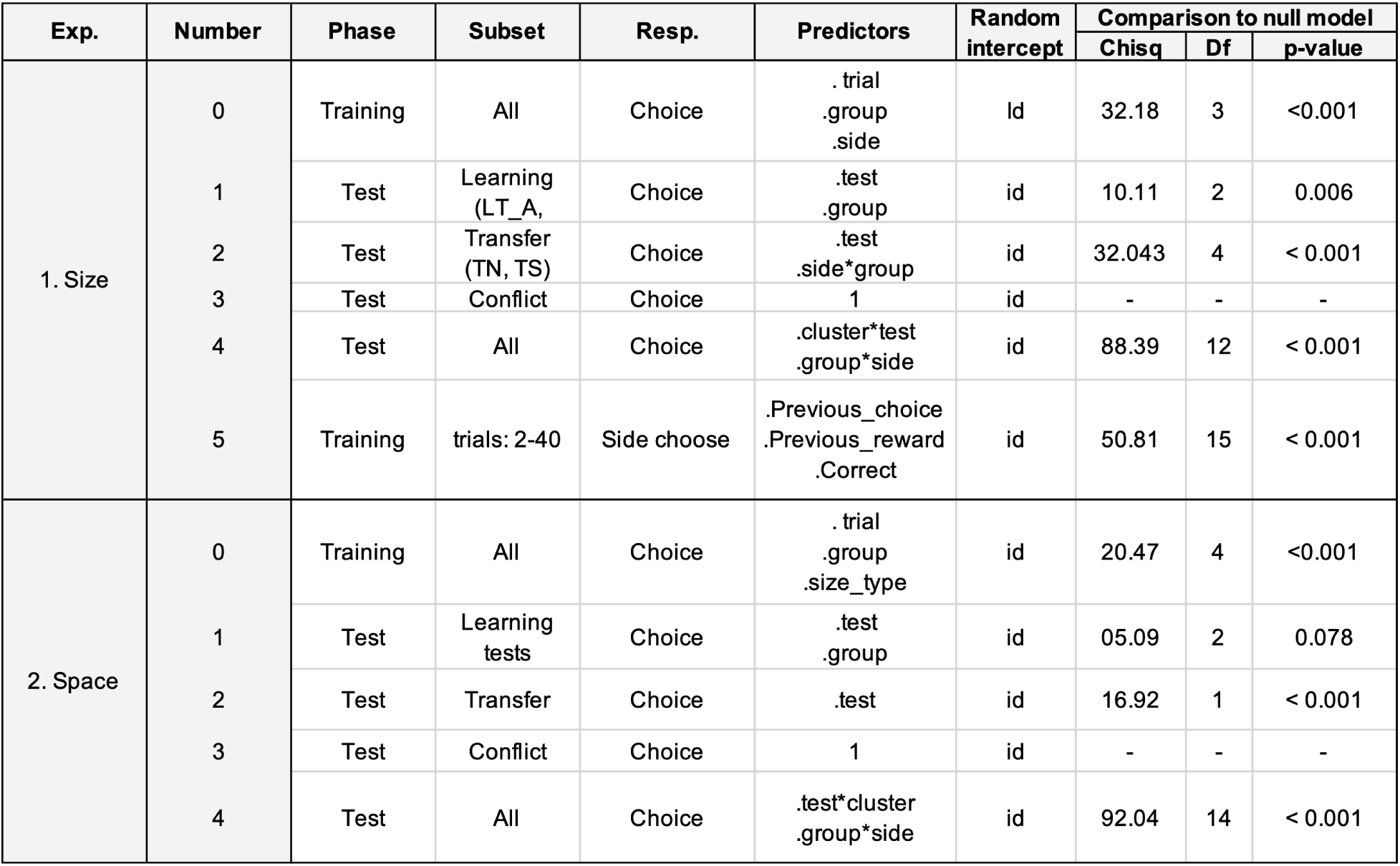
Summary of statistical models. Each line corresponds to one specific model. Training phase predictors: trial (1-40) indexes each learning trial; space_type defines the Space control applied to training stimuli of Experiment Size (A: convex hull; B: density; C: inter-distance); size_type defines the Size control applied to training stimuli of Experiment Space (A: area; B: perimeter; C: item size); group indicates the rewarded numerosity during training (2:CS+ or 4:CS+); side specifies the arm of the Y-maze where the correct stimulus was displayed (L or R). Testing phase predictors: test identifies the type of test (Experiment Size: LT_A, LT_C, TN, TS, CNS; Experiment Space: LT_B, LT_C, TN, TSp, CNS); test_order_all (1-5) gives the sequential position of each test in the testing phase; testing_set identifies the specific stimulus pair presented; cluster denotes the behavioral subgroup identified via hierarchical clustering (i: numerical bias; ii: generalist). Heuristic and rule-learning predictors: chosen(t–1) indicates the side chosen on the previous trial (L or R); Reward(t–1) indicates the side where sucrose was obtained on the previous trial (L or R); Correct(t) indicates the side where the correct stimulus appeared on the current trial (L or R). P-values correspond to full vs. null model comparisons. df: degrees of freedom; Exp.: experiment, id.: individual; Resp.: response variable.

| Exp. | Number | Phase | Subset | Resp. | Predictors | Random intercept | Comparison to null model |  |  |
| --- | --- | --- | --- | --- | --- | --- | --- | --- | --- |
|  |  |  |  |  |  |  | Chisq | Df | p-value |
| 1. Size | 0 | Training | All | Choice | .trial<br>.group<br>.side | id | 32.18 | 3 | <0.001 |
|  | 1 | Test | Learning (LT_A, | Choice | .test<br>.group | id | 10.11 | 2 | 0.006 |
|  | 2 | Test | Transfer (TN, TS) | Choice | .test<br>.side*group | id | 32.043 | 4 | < 0.001 |
|  | 3 | Test | Conflict | Choice | 1 | id | - | - | - |
|  | 4 | Test | All | Choice | .cluster*test<br>.group*side | id | 88.39 | 12 | < 0.001 |
|  | 5 | Training | trials: 2-40 | Side choose | .Previous_choice<br>.Previous_reward<br>.Correct | id | 50.81 | 15 | < 0.001 |
| 2. Space | 0 | Training | All | Choice | .trial<br>.group<br>.size_type | id | 20.47 | 4 | <0.001 |
|  | 1 | Test | Learning tests | Choice | .test<br>.group | id | 05.09 | 2 | 0.078 |
|  | 2 | Test | Transfer | Choice | .test | id | 16.92 | 1 | < 0.001 |
|  | 3 | Test | Conflict | Choice | 1 | id | - | - | - |
|  | 4 | Test | All | Choice | .test*cluster<br>.group*side | id | 92.04 | 14 | < 0.001 |

**Fig S1.**
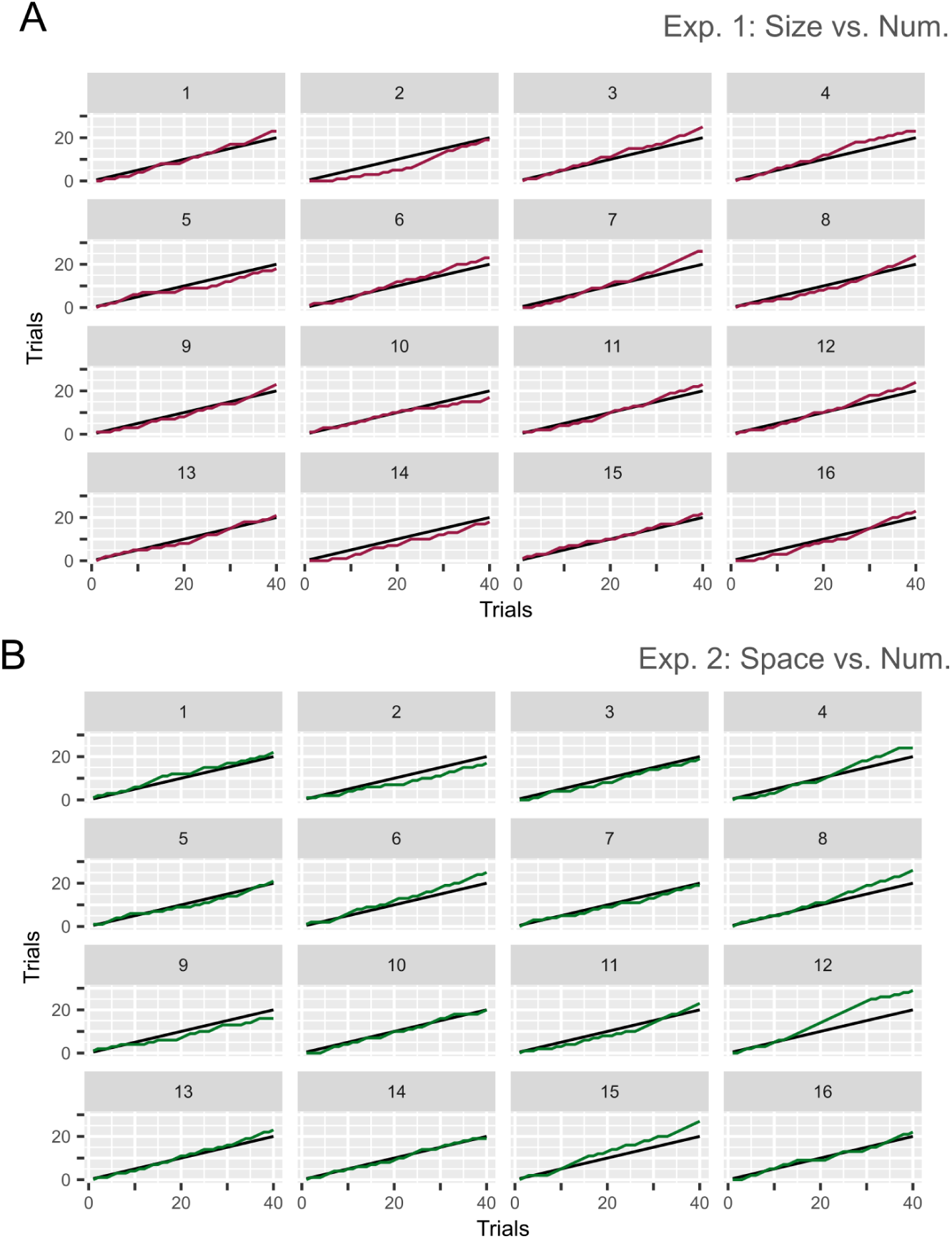
Individual learning curves of honeybees in Experiment Size and 2: A) Cumulative correct choices of individual bees in Experiment Size. B) Cumulative correct choices of individual bees in Experiment Space. Each panel corresponds to a single bee identified by a number from 1 to 16. The black line corresponds to the evolution of cumulative correct choices according to random choices.

**Fig S2.**
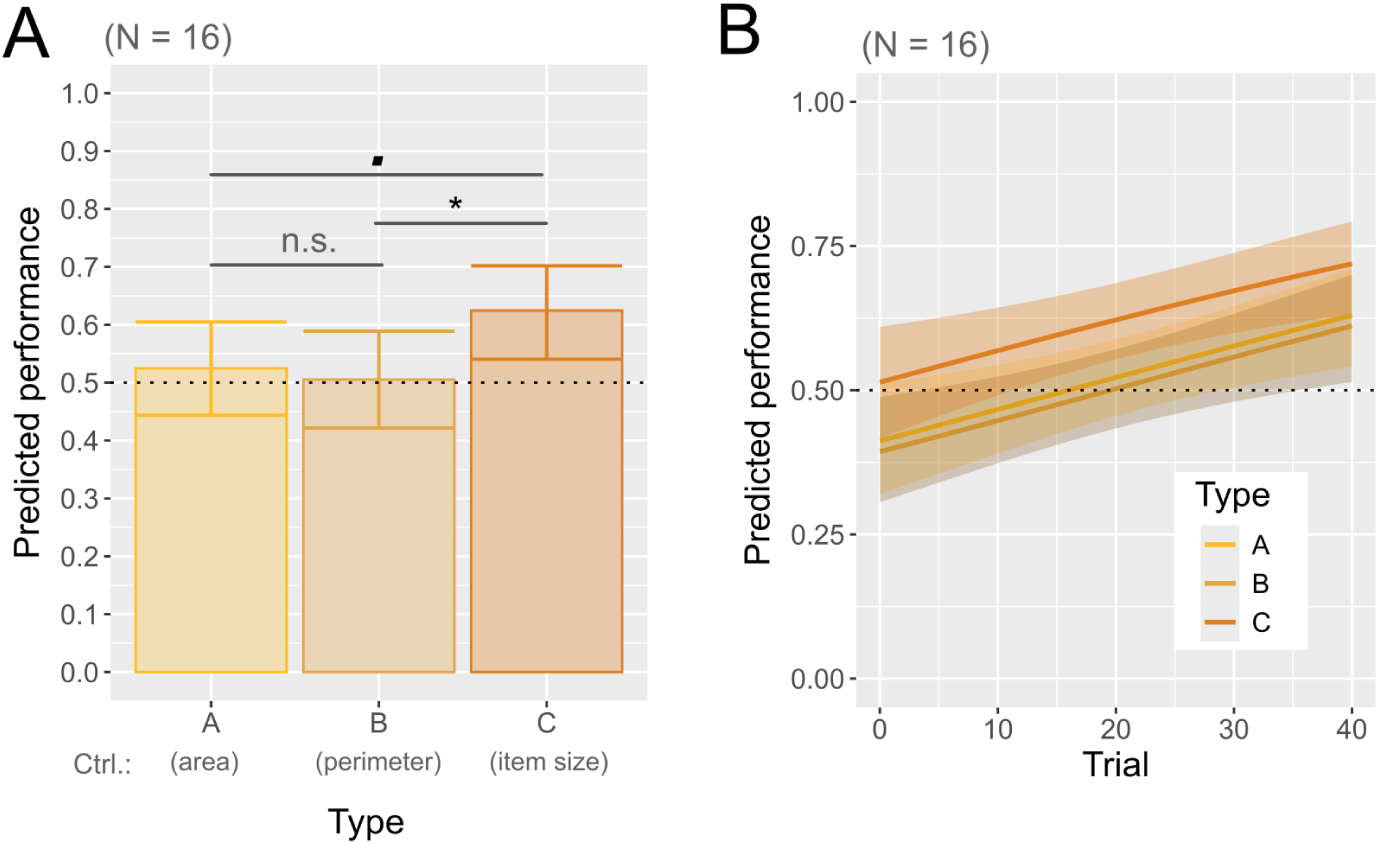
Size control during training phase in Experiment Space: A) Predicted mean performance of bees for each stimulus type. Bars denote predicted mean. Error bars show 95% CIs. B) Evolution of the effect of Size-type control on the probability to make a correct choice. Thick lines denote predicted mean. Shaded areas show 95% confidence intervals. The black dotted line indicates chance level. (***): p ≤ 0.001, (**): p ≤ 0.01, (*): p ≤ 0.05, (■): p ≤ 0.1, (n.s.): p ě 0.1.

**Fig S3.**
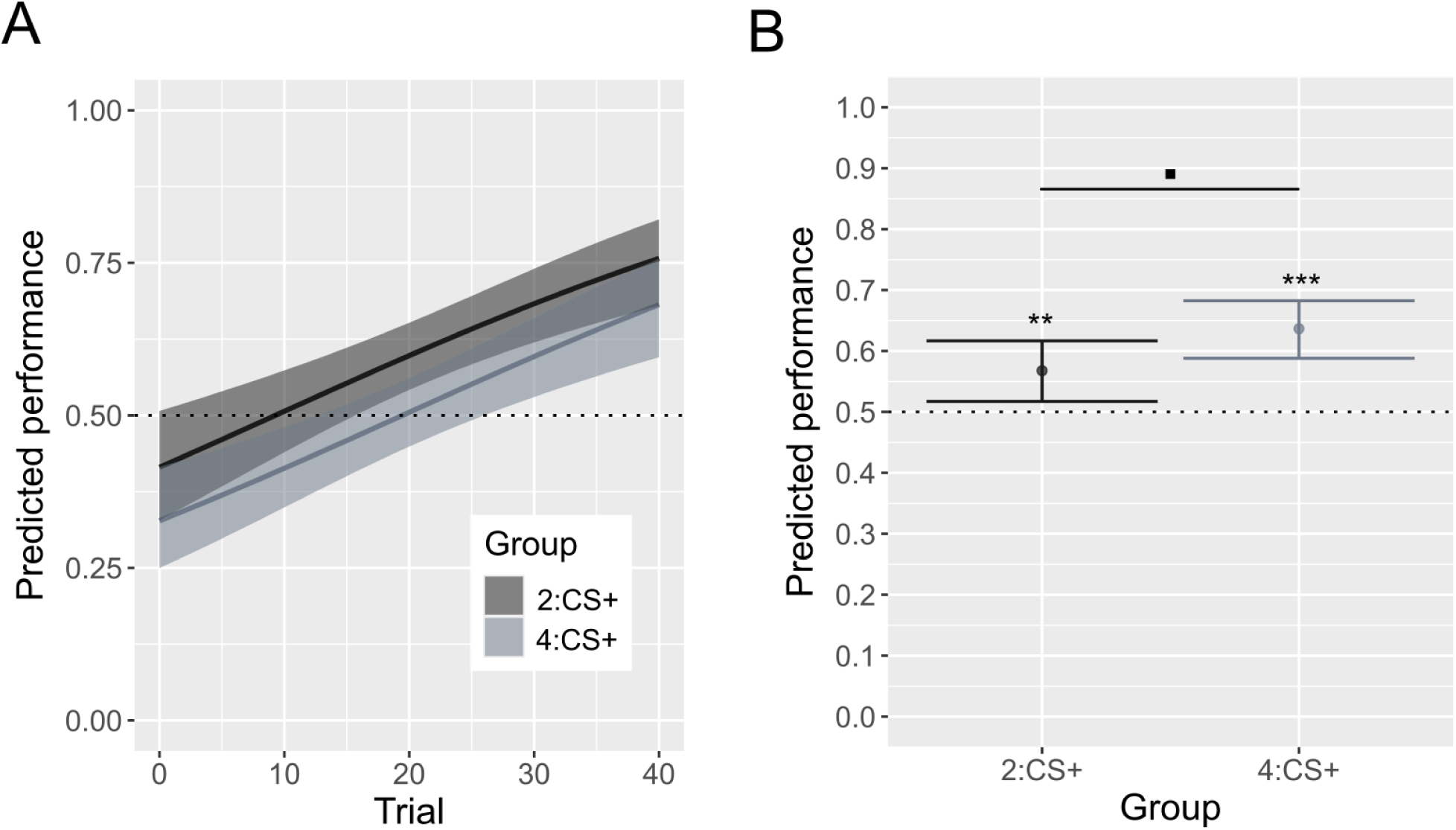
Effect of training group during the training and testing phases of Experiment Size. A) Predicted performance of bees according to their training group across the training phase (2:CS+: N = 8, n = 320; 4:CS+: N = 8, n = 320). B) Predicted performance of bees from both training groups in the learning tests (2:CS+: N = 8, n = 724; 4:CS+: N = 8, n = 735). Dots represent the predicted mean performance across learning tests, with error bars showing 95% confidence intervals. The black dotted line indicates chance level. (***): p ≤ 0.001, (**): p ≤ 0.01, (*): p ≤ 0.05, (■): p ă 0.1, (n.s.): p ě 0.1.

**Fig S4.**
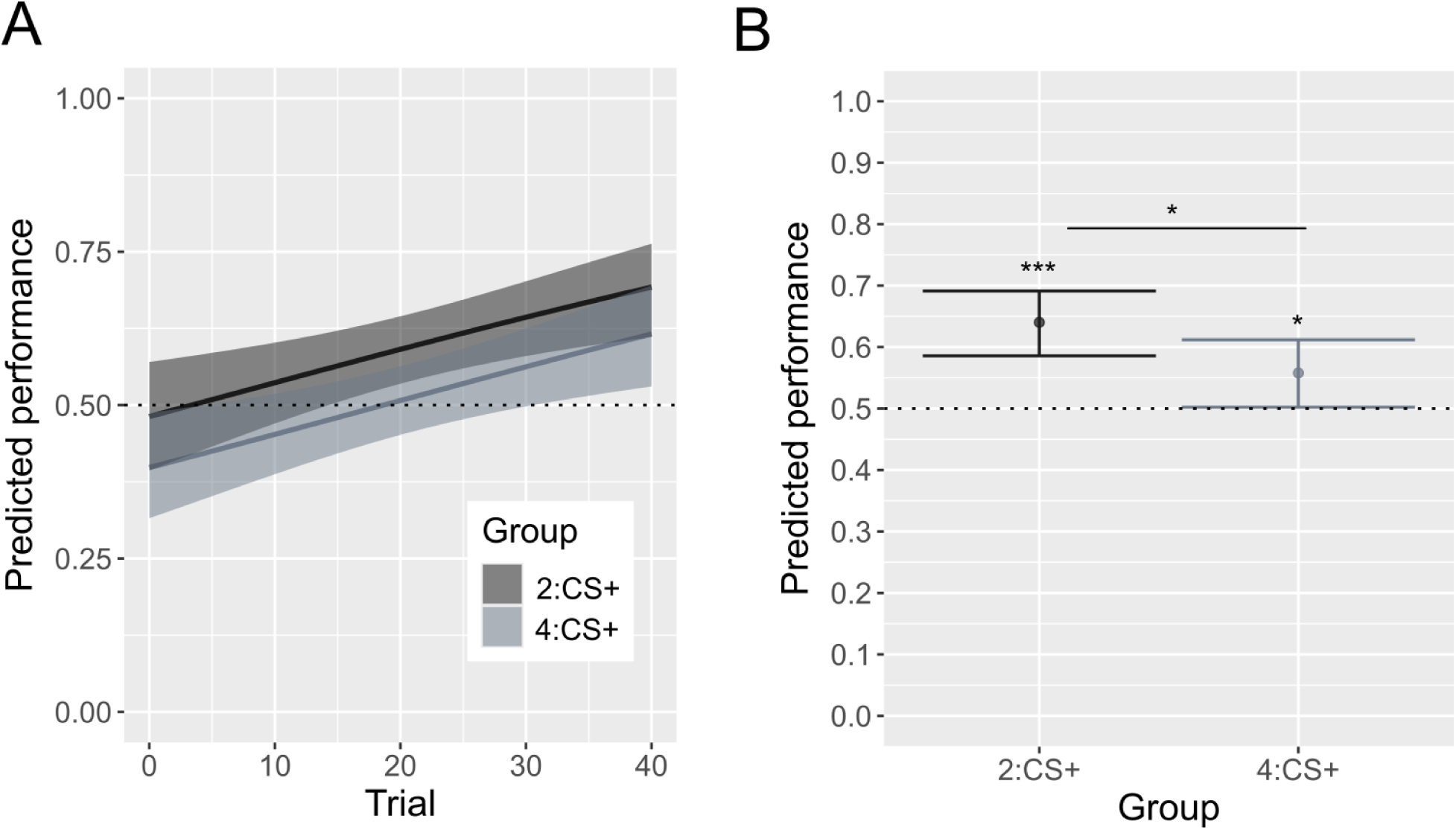
Effect of group on the training and testing phases of Experiment Space. A) Predicted performance of bees from each training group across the training phase (2:CS+: N = 8, n = 320; 4:CS+: N = 8, n = 320). B) Predicted performance of bees from both training groups in the learning tests (2:CS+: N = 8, n = 599; 4:CS+: N = 8, n = 628). Bars show the mean proportion of correct choices for each group across both tests. Error bars indicate 95% CIs. The black dotted line represents chance-level performance. Significance codes: (***) p ≤ 0.001, (**) p ≤ 0.01, (*) p ≤ 0.05, (■) p ă 0.1, (n.s.) p ě 0.1.

**Fig S5.**
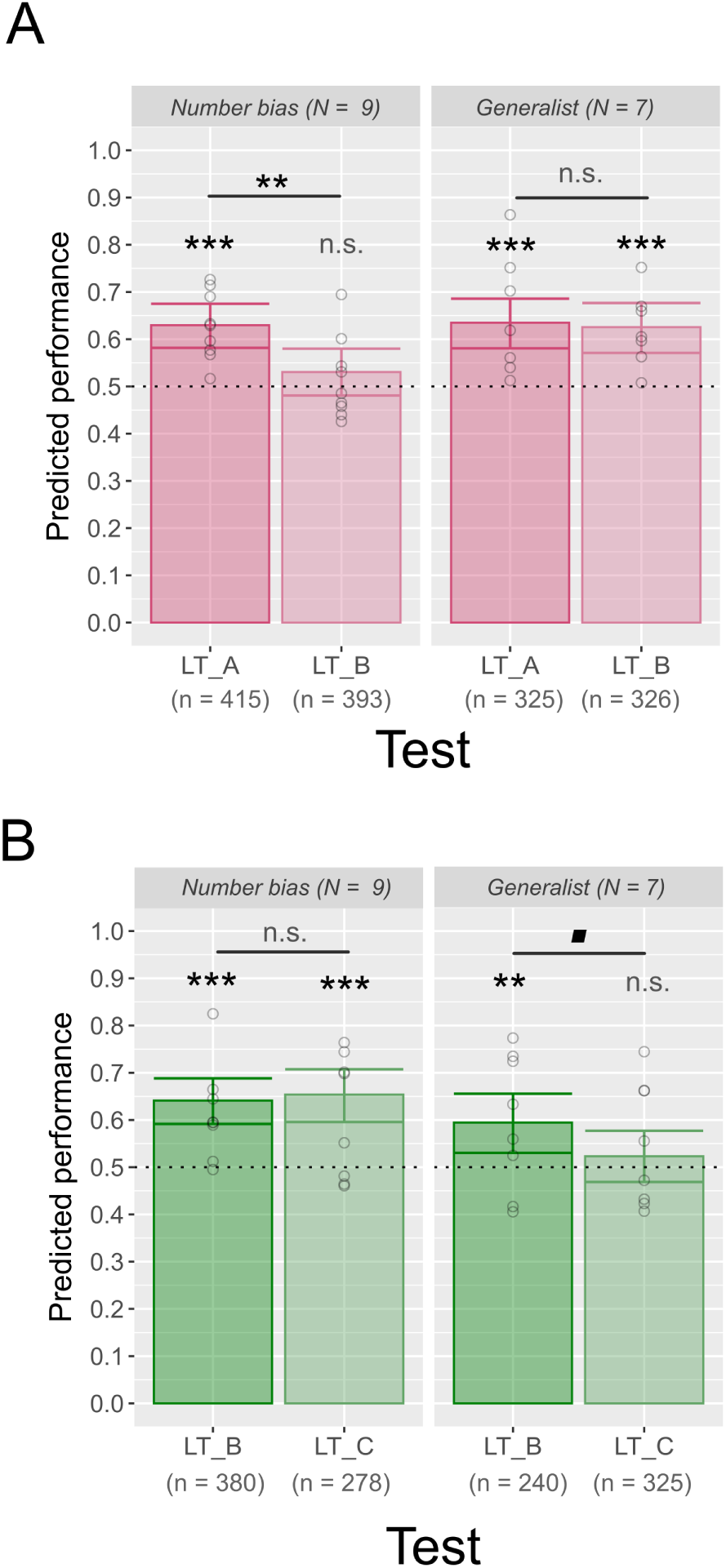
Learning test of bees according to the cluster. Experiment Size: A) Predicted performance in learning for bees in the first “number bias” and second “generalist” cluster in Experiment Size. Data and significance for come from model 1.4. B) Predicted performance in learning for bees in the first “number bias” and second “generalist” cluster in Experiment Space. Data and significance come from model 2.4. Each bar shows the predicted mean, and error bars denote 95% CIs. Each point represents the performance of one bee in a single test bout (raw data). The dotted horizontal line indicates chance level (0.5). (***): p ≤ 0.001, (**): p ≤ 0.01, (*): p ≤ 0.05, (■): p ≤ 0.1, (n.s.): p > 0.1. LT: learning test; N: number of individuals; n: number of observations.

**Fig S6.**
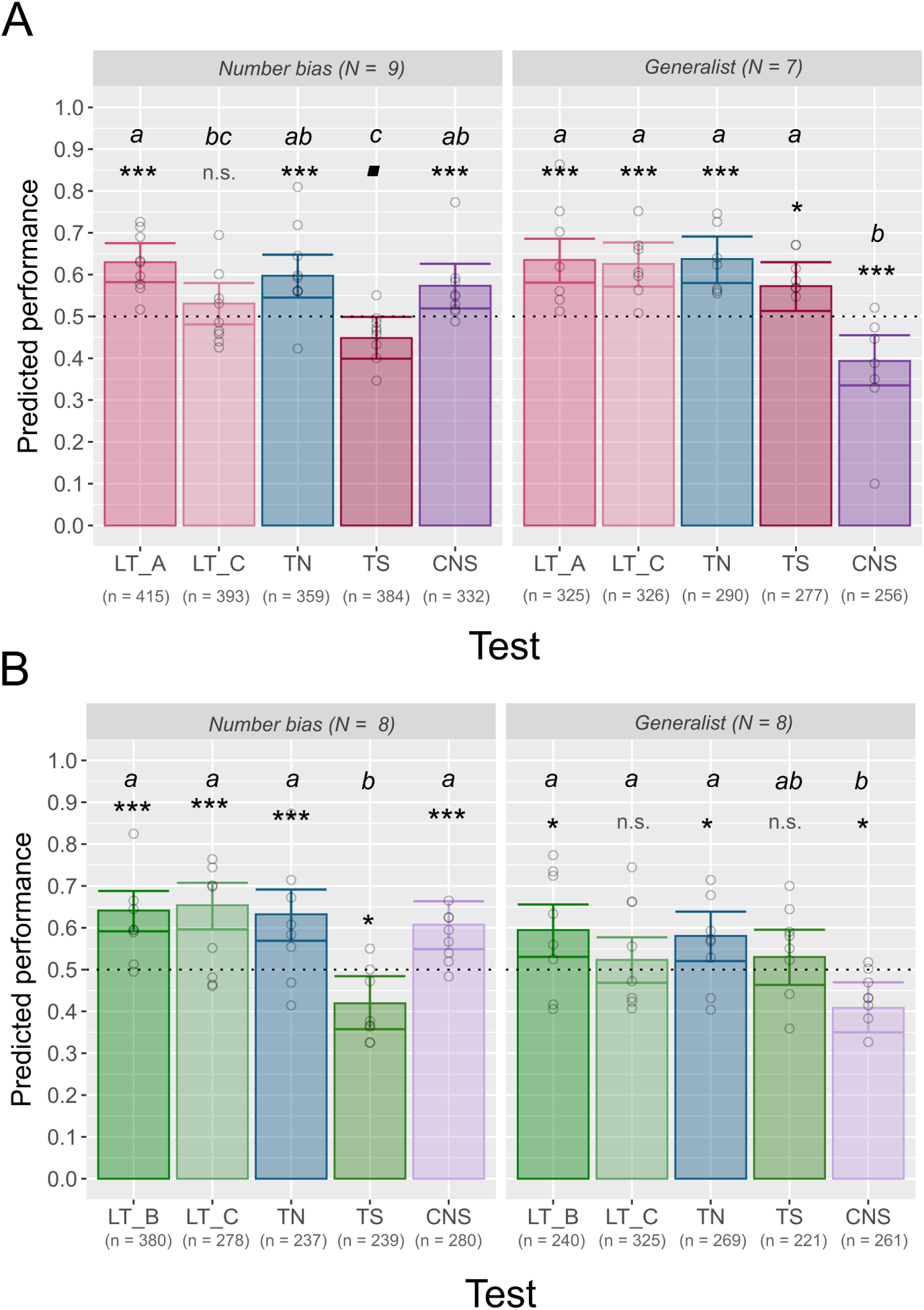
Test performance of bees according to the cluster. A) Predicted performance in learning, transfer, and conflict tests for bees in the first “number bias” and second “generalist” cluster in Experiment Size. Data and significance come from model 1.4. B) Predicted performance in learning, transfer and conflict tests for bees in the first “number bias” and second “generalist” cluster in Experiment Space. Data and significance come from model 2.4. Each bar shows the predicted mean, and error bars denote 95% CIs. Each point represents the performance of one bee in a single test bout (raw data). The dotted horizontal line indicates chance level (0.5). Different letters indicate significant pairwise differences between groups (holm-adjusted, p < 0.05). (***): p ≤ 0.001, (**): p ≤ 0.01, (*): p ≤ 0.05, (■): p ≤ 0.1, (n.s.): p > 0.1. LT: learning test; N: number of individuals; n: number of observations.

**Fig S7.**
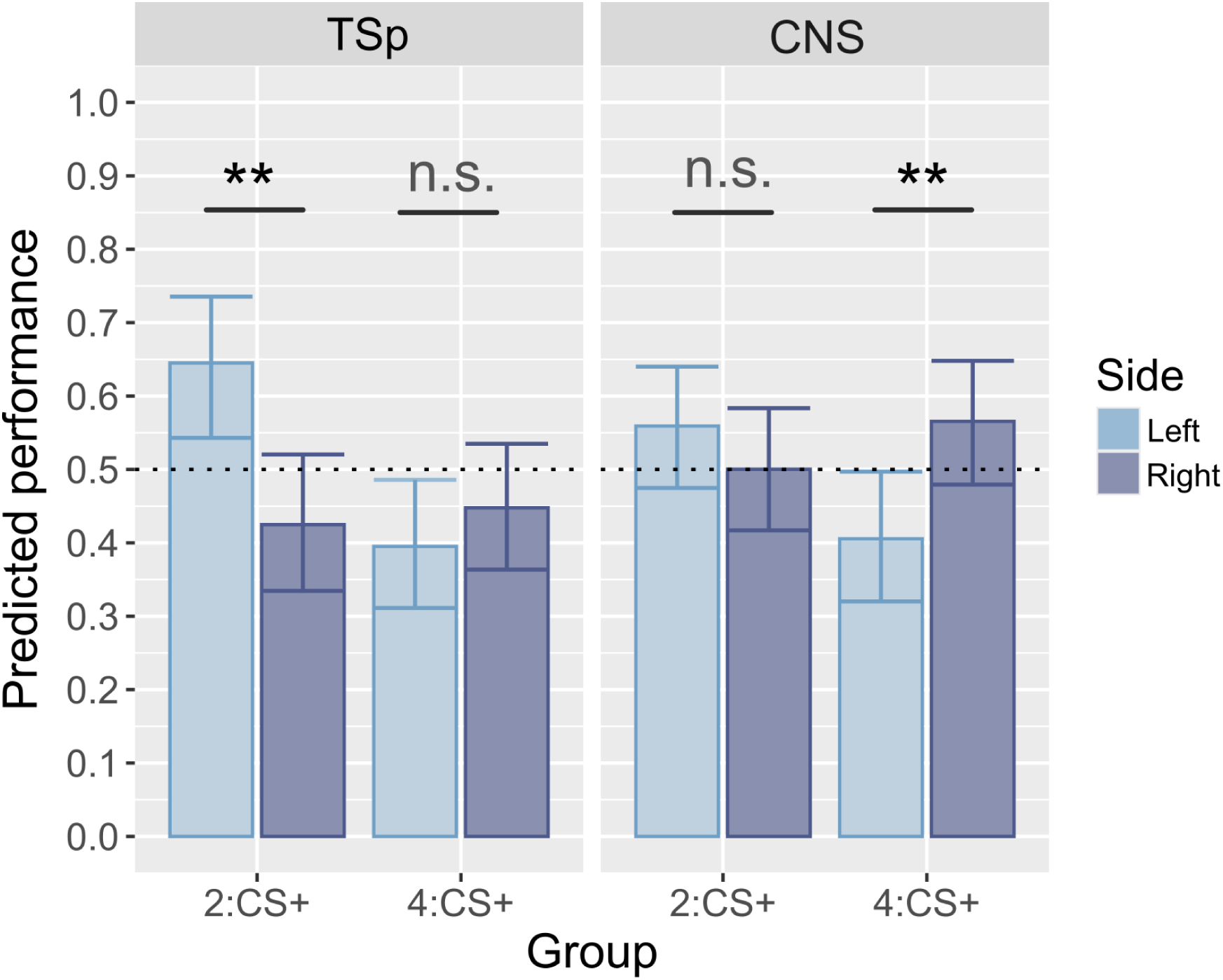
Left-to-right congruency effect in Experiment Space. Predicted performance in learning, transfer and conflict tests. Data and significance come from model 2.5. Each small horizontal bar indicates the predicted mean, and error bars denote 95% CIs. The dotted horizontal line indicates chance level (0.5). (**): p ≤ 0.01, (n.s.): p > 0.1. CNS: conflict test; LT: learning test; N: number of individuals; n: number of observations; Num.: numerosity; TN: numerical transfer; TSp: Space transfer.

**Fig S8.**
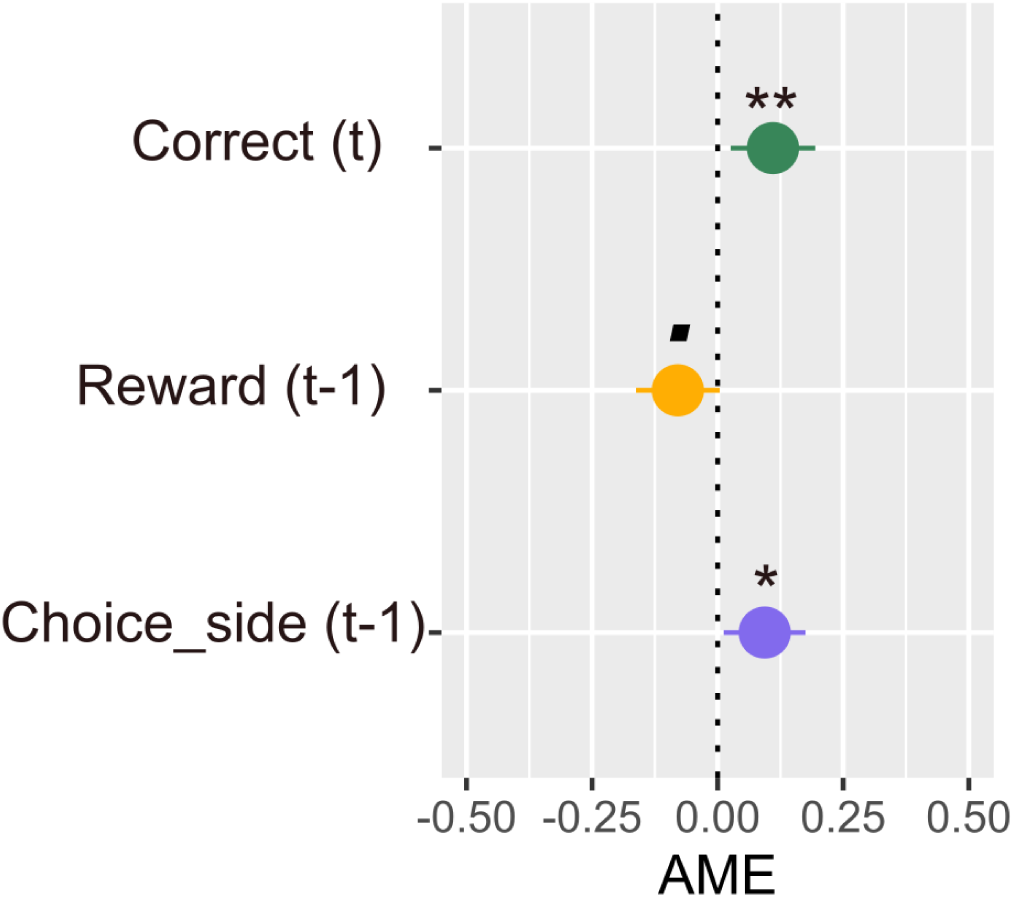
From heuristics to rule learning: navigating a Y-maze in Experiment Space. Average marginal effects of three predictors on the side chosen by bees in Experiment Space. Reward(t–1) indicates the side where the reward was obtained on the previous trial; Choice_side(t–1) the side chosen on the previous trial; Correctptq the side where the correct stimulus was displayed. The vertical dashed line represents zero effect, corresponding to chance (no influence on side choice). (**): p ≤ 0.01, (*): p ≤ 0.05, (■): p ≤ 0.1, (n.s.): p > 0.1. AME: Average marginal effect.

## References

1. Abramson, J. Z., Hernández-Lloreda, V., Call, J., & Colmenares, F. (2011). Relative quantity judgments in South American sea lions (*Otaria flavescens*). Animal cognition, 14 (5), 695–706. 10.1007/s10071-011-0404-7

2. Abramson, J. Z., Hernández-Lloreda, V., Call, J., & Colmenares, F. (2013). Relative quantity judgments in the beluga whale (*Delphinapterus leucas*) and the bottlenose dolphin (*Tursiops truncatus*). Behavioural Processes, 96, 11–19. 10.1016/j.beproc.2013.02.006

3. Adam, E., Zanon, M., Messina, A., & Vallortigara, G. (2024). Looks like home: Numerosity, but not spatial frequency guides preference in zebrafish larvae (Danio rerio). Animal Cognition, 27 (1), 53. 10.1007/s10071-024-01888-0

4. Adriano, A., Girelli, L., & Rinaldi, L. (2021). Non-symbolic numerosity encoding escapes spatial frequency equalization. Psychological Research, 85 (8), 3061–3074. 10.1007/s00426-020-01458-2

5. Agrillo, C. (2015). Numerical and arithmetic abilities in non-primate species. In The Oxford handbook of numerical cognition (pp. 214–236). Oxford University Press Oxford, UK.

6. Agrillo, C., & Bisazza, A. (2018). Understanding the origin of number sense: A review of fish studies. Philosophical Transactions of the Royal Society B: Biological Sciences, 373 (1740), 20160511. 10.1098/rstb.2016.0511

7. Agrillo, C., Dadda, M., Serena, G., & Bisazza, A. (2008). Do fish count? Spontaneous discrimination of quantity in female mosquitofish. Animal Cognition, 11 (3), 495–503. 10.1007/s10071-008-0140-9

8. Agrillo, C., Dadda, M., Serena, G., & Bisazza, A. (2009). Use of number by fish. PloS one, 4 (3), e4786. 10.1371/journal.pone.0004786

9. Agrillo, C., Piffer, L., & Bisazza, A. (2010). Large number discrimination by mosquitofish. PloS one, 5 (12), e15232. 10.1371/journal.pone.0015232

10. Agrillo, C., Piffer, L., & Bisazza, A. (2011). Number versus continuous quantity in numerosity judgments by fish. Cognition, 119 (2), 281–287. 10.1016/j.cognition.2010.10.022

11. Aın, S. A., Giret, N., Grand, M., Kreutzer, M., & Bovet, D. (2009). The discrimination of discrete and continuous amounts in African grey parrots (*Psittacus erithacus*). Animal cognition, 12 (1), 145–154. 10.1007/s10071-008-0178-8

12. Anobile, G., Cicchini, G. M., & Burr, D. C. (2014). Separate mechanisms for perception of numerosity and density. Psychological science, 25 (1), 265–270. 10.1177/0956797613501520

13. Aulet, L. S., & Lourenco, S. F. (2023). No intrinsic number bias: Evaluating the role of perceptual discriminability in magnitude categorization. Developmental Science, 26 (2), e13305. 10.1111/desc.13305

14. Avargues-Weber, A., Dyer, A. G., Ferrah, N., & Giurfa, M. (2015). The forest or the trees: Preference for global over local image processing is reversed by prior experience in honeybees. Proceedings of the Royal Society B: Biological Sciences, 282 (1799), 20142384. 10.1098/rspb.2014.2384

15. Avargues-Weber, A., & Giurfa, M. (2015). Conceptual learning by miniature brains. 10.1098/rspb.2013.1907

16. Avarguès-Weber, A., Portelli, G., Benard, J., Dyer, A., & Giurfa, M. (2010). Configural processing enables discrimination and categorization of face-like stimuli in honeybees. Journal of Experimental Biology, 213 (4), 593–601. 10.1242/jeb.039263

17. Avarguès-Weber, A., d’Amaro, D., Metzler, M., & Dyer, A. G. (2014). Conceptualization of relative size by honeybees. Frontiers in Behavioral Neuroscience, 8, 80. 10.3389/fnbeh.2014.00080

18. Avarguès-Weber, A., de Brito Sanchez, M. G., Giurfa, M., & Dyer, A. G. (2010). Aversive reinforcement improves visual discrimination learning in free-flying honeybees. PLoS One, 5 (10), e15370. 10.1371/journal.pone.0015370

19. Avarguès-Weber, A., Dyer, A. G., Combe, M., & Giurfa, M. (2012). Simultaneous mastering of two abstract concepts by the miniature brain of bees. Proceedings of the National Academy of Sciences, 109 (19), 7481–7486. 10.1073/pnas.1202576109

20. Avarguès-Weber, A., Dyer, A. G., & Giurfa, M. (2011). Conceptualization of above and below relationships by an insect. Proceedings of the Royal Society B: Biological Sciences, 278 (1707), 898–905. 10.1098/rspb.2010.1891

21. Bánszegi, O., Urrutia, A., Szenczi, P., & Hudson, R. (2016). More or less: Spontaneous quantity discrimination in the domestic cat. Animal Cognition, 19 (5), 879–888. 10.1007/s10071-016-0985-2

22. Baracchi, D., Rigosi, E., de Brito Sanchez, G., & Giurfa, M. (2018). Lateralization of sucrose responsiveness and non-associative learning in honeybees. Frontiers in Psychology, 9, 425. 10.3389/fpsyg.2018.00425

23. Bengochea, M., Sitt, J. D., Izard, V., Preat, T., Cohen, L., & Hassan, B. A. (2023). Numerical discrimination in *Drosophila melanogaster*. Cell Reports, 42 (7), 112772. 10.1016/j.celrep.2023.112772

24. Benson-Amram, S., Heinen, V. K., Dryer, S. L., & Holekamp, K. E. (2011). Numerical assessment and individual call discrimination by wild spotted hyaenas, <I>Crocuta crocuta <I>. Animal Behaviour, 82 (4), 743–752. 10.1016/j.anbehav.2011.07.004

25. Beran, M. J., French, K., Smith, T. R., & Parrish, A. E. (2019). Limited evidence of number-space mapping in rhesus monkeys (<I> Macaca mulatta <I>) and capuchin monkeys (<I> Sapajus apella <I>). Journal of Comparative Psychology, 133 (3), 281. 10.1037/com0000177

26. Bisazza, A., & Gatto, E. (2021). Continuous versus discrete quantity discrimination in dune snail (Mollusca: Gastropoda) seeking thermal refuges. Scientific Reports, 11 (1), 3757. 10.1038/s41598-021-82249-6

27. Bisazza, A., Tagliapietra, C., Bertolucci, C., Foà, A., & Agrillo, C. (2014). Non-visual numerical discrimination in a blind cavefish (<I> Phreatichthys andruzzii <I>). Journal of Experimental Biology, 217 (11), 1902–1909. 10.1242/jeb.101683

28. Bortot, M., Agrillo, C., Avarguès-Weber, A., Bisazza, A., Miletto Petrazzini, M. E., & Giurfa, M. (2019). Honeybees use absolute rather than relative numerosity in number discrimination. Biology Letters, 15 (6), 20190138. 10.1098/rsbl.2019.0138

29. Bortot, M., Regolin, L., & Vallortigara, G. (2021). A sense of number in invertebrates. Biochemical and Biophysical Research Communications, 564, 37–42. 10.1016/j.bbrc.2020.11.039

30. Bortot, M., Stancher, G., & Vallortigara, G. (2020). Transfer from number to size reveals abstract coding of magnitude in honeybees. iScience, 23 (5), 101122. 10.1016/j.isci.2020.101122

31. Bortot, M., & Vallortigara, G. (2023). Transfer from continuous to discrete quantities in honeybees. iScience, 26 (10), 108035. 10.1016/j.isci.2023.108035

32. Bulf, H., De Hevia, M. D., & Macchi Cassia, V. (2016). Small on the left, large on the right: Numbers orient visual attention onto space in preverbal infants. Developmental Science, 19 (3), 394–401. 10.1111/desc.12315

33. Bulut, M., Hepdarcan, I., Palaz, E., Çetinkaya, H., & Dural, S. (2023). No SNARC effect among left-to-right readers: Evidence from a Turkish sample. Advances in Cognitive Psychology. 10.5709/acp-0394-x

34. Cantlon, J. F., & Brannon, E. M. (2006). Shared system for ordering small and large numbers in monkeys and humans. Psychological science, 17 (5), 401–406. 10.1111/j.1467-9280.2006.01719.x

35. Carazo, P., Fernández-Perea, R., & Font, E. (2012). Quantity estimation based on numerical cues in the mealworm beetle (*Tenebrio molitor*). Frontiers in Psychology, 3, 502. 10.3389/fpsyg.2012.00502

36. Casanova, A., Rodríguez, C., Ferreira, R. A., & Miranda, I. (2025). Exploring the association between inhibitory control and mathematical performance in early childhood: A systematic review. Applied Neuropsychology: Child, 1–23. 10.1080/21622965.2025.2552198

37. Chittka, L., & Geiger, K. (1995). Can honey bees count landmarks? Animal Behaviour, 49 (1), 159–164. 10.1016/0003-3472(95)80163-4

38. Cicchini, G. M., Anobile, G., & Burr, D. C. (2016). Spontaneous perception of numerosity in humans. Nature communications, 7 (1), 12536. 10.1038/ncomms12536

39. Claeys, E., Kerjean, E., & Loubes, J.-M. (2025). Buzz, choose, forget: A meta-bandit framework for bee-like decision making. arXiv preprint arXiv:2510.16462.

40. Clayton, S., Gilmore, C., & Inglis, M. (2015). Dot comparison stimuli are not all alike: The effect of different visual controls on ANS measurement. Acta Psychologica, 161, 177–184. 10.1016/j.actpsy.2015.09.007

41. Cochran, W. T., Cooley, J. W., Favin, D. L., Helms, H. D., Kaenel, R. A., Lang, W. W., Maling, G. C., Nelson, D. E., Rader, C. M., & Welch, P. D. (1967). What is the fast Fourier transform? Proceedings of the IEEE, 55 (10), 1664–1674. 10.1109/PROC.1967.5957

42. Cook, C. N., Mosqueiro, T., Brent, C. S., Ozturk, C., Gadau, J., Pinter-Wollman, N., & Smith, B. H. (2019). Individual differences in learning and biogenic amine levels influence the behavioural division between foraging honeybee scouts and recruits. Journal of Animal Ecology, 88 (2), 236–246. 10.1111/1365-2656.12911

43. Cooper, T. L., Pardo-Sanchez, J., Sosnowski, M. J., Rodriguez, S. T., Spencer, M. S., Brosnan, S. F., & Mendelson III, J. R. (2024). How to tell more is more: Quantity discrimination in eastern box turtles (emydidae: <I> Terrapene carolina <I>). Journal of Herpetology, 58 (1), 1–15. 10.1670/23-018

44. Cox, L., & Montrose, V. T. (2016). Quantity discrimination in domestic rats, *Rattus norvegicus*. Animals, 6 (8), 46. 10.3390/ani6080046

45. Dacke, M., & Srinivasan, M. V. (2008). Evidence for counting in insects. Animal Cognition, 11, 683–689. 10.1007/s10071-008-0159-y

46. Dantzig, T. (1931). Number; the language of science. MacMillan, 20.

47. Davidson, R. J. (2004). Well-being and affective style: Neural substrates and biobehavioural correlates. Philosophical Transactions of the Royal Society of London. Series B: Biological Sciences, 359 (1449), 1395–1411. 10.1098/rstb.2004.1510

48. Davis, H., & Memmott, J. (1982). Counting behavior in animals: A critical evaluation. Psychological Bulletin, 92 (3), 547. 10.1037/0033-2909.92.3.547

49. Davis, H., & Pérusse, R. (1988). Numerical competence in animals: Definitional issues, current evidence, and a new research agenda. Behavioral and Brain Sciences, 11 (4), 561–579. 10.1017/S0140525X00053437

50. de Hevia, M. D., Veggiotti, L., Streri, A., & Bonn, C. D. (2017). At birth, humans associate “few” with left and “many” with right. Current Biology, 27 (24), 3879–3884. 10.1016/j.cub.2017.11.024

51. De Hevia, M. D., Girelli, L., Addabbo, M., & Macchi Cassia, V. (2014). Human Infants’ Preference for Left-to-Right Oriented Increasing Numerical Sequences (E. Kroesbergen, Ed.). PLoS ONE, 9 (5), e96412. 10.1371/journal.pone.0096412

52. Dehaene, S. (1997). The number sense: How the mind creates mathematics.

53. Dehaene, S., Bossini, S., & Giraux, P. (1993). The mental representation of parity and number magnitude. Journal of experimental psychology: General, 122 (3), 371. 10.1037//0096-3445.122.3.371

54. DeWind, N. K., Adams, G. K., Platt, M. L., & Brannon, E. M. (2015). Modeling the approximate number system to quantify the contribution of visual stimulus features. Cognition, 142, 247–265. 10.1016/j.cognition.2015.05.016

55. Diaz, F., & Wasserman, E. A. (2023). The role of numerical and nonnumerical magnitudes in pigeons’ conditional discrimination behavior. Journal of Experimental Psychology: Animal Learning and Cognition, 49 (4), 253. 10.1037/xan0000368

56. Diaz, F., & Wasserman, E. A. (n.d.). Nonnumerical stimuli exert surprisingly strong behavioral control in an unconstrained numerical discrimination learning task. Available at SSRN 5175053. 10.2139/ssrn.5175053

57. Ditz, H. M., & Nieder, A. (2016). Numerosity representations in crows obey the Weber-Fechner law. Proceedings of the Royal Society B: Biological Sciences, 283 (1827), 20160083. 10.1098/rspb.2016.0083

58. Dos Santos, C. F. (2022). Re-establishing the distinction between numerosity, numerousness, and number in numerical cognition. Philosophical Psychology, 35 (8), 1152–1180. 10.1080/09515089.2022.2029387

59. Drucker, C. B., & Brannon, E. M. (2014). Rhesus monkeys (*Macaca mulatta*) map number onto space. Cognition, 132 (1), 57–67. 10.1016/j.cognition.2014.03.011

60. Eccher, E., Josserand, M., Caparos, S., Boissin, E., Buiatti, M., Piazza, M., & Vallortigara, G. (2025). A left-to-right bias in number-space mapping across ages and cultures. Nature Communications, 16 (1), 495. 10.1038/s41467-024-55685-x

61. Eckert, J., Bohn, M., & Spaethe, J. (2022). Does quantity matter to a stingless bee? Animal Cognition, 25 (3), 617–629. 10.1007/s10071-021-01581-6

62. Emmerton, J. (1998). Numerosity differences and effects of stimulus density on pigeons’ discrimination performance. Animal Learning & Behavior, 26 (3), 243–256. 10.3758/BF03199218

63. Estes, K. W. (1976). Nonverbal discrimination of more and fewer elements by children. Journal of Experimental Child Psychology, 21 (3), 393–405. 10.1016/0022-0965(76)90069-2

64. Ferreira, C. H., & Moita, M. A. (2020). Behavioral and neuronal underpinnings of safety in numbers in fruit flies. Nature communications, 11 (1), 4182. 10.1038/s41467-020-17856-4

65. Ferrigno, S., Jara-Ettinger, J., Piantadosi, S. T., & Cantlon, J. F. (2017). Universal and uniquely human factors in spontaneous number perception. Nature communications, 8 (1), 13968. 10.1038/ncomms13968

66. Fischer, M. H., Castel, A. D., Dodd, M. D., & Pratt, J. (2003). Perceiving numbers causes spatial shifts of attention. Nature neuroscience, 6 (6), 555–556. 10.1038/nn1066

67. Frasnelli, E., Haase, A., Rigosi, E., Anfora, G., Rogers, L. J., & Vallortigara, G. (2014). The bee as a model to investigate brain and behavioural asymmetries. Insects, 5 (1), 120–138. 10.3390/insects5010120

68. Garland, A., Low, J., & Burns, K. C. (2012). Large quantity discrimination by North Island robins (<I> Petroica longipes <I>). Animal cognition, 15 (6), 1129–1140. 10.1007/s10071-012-0537-3

69. Gatto, E., Loukola, O. J., & Agrillo, C. (2022). Quantitative abilities of invertebrates: A methodological review. Animal Cognition, 25 (1), 5–19. 10.1007/s10071-021-01529-w

70. Gebuis, T., Cohen Kadosh, R., & Gevers, W. (2016). Sensory-integration system rather than approximate number system underlies numerosity processing: A critical review. Acta Psychologica, 171, 17–35. 10.1016/j.actpsy.2016.09.003

71. Gebuis, T., & Reynvoet, B. (2011). Generating nonsymbolic number stimuli. Behavior research methods, 43 (4), 981–986. 10.3758/s13428-011-0097-5

72. Giurfa, M. (2019). An Insect’s Sense of Number. Trends in Cognitive Sciences, 23 (9), 720–722. 10.1016/j.tics.2019.06.010

73. Giurfa, M. (2022). Pollinator cognition: Framing bee memories in an ecological context. Current Biology, 32 (19), R1015–R1018. 10.1016/j.cub.2022.08.043

74. Giurfa, M., Marcout, C., Hilpert, P., Thevenot, C., & Rugani, R. (2022). An insect brain organizes numbers on a left-to-right mental number line. Proceedings of the National Academy of Sciences, 119 (44), e2203584119. 10.1073/pnas.2203584119

75. Giurfa, M., Vorobyev, M., Kevan, P., & Menzel, R. (1996). Detection of coloured stimuli by honeybees: Minimum visual angles and receptor specific contrasts. Journal of Comparative Physiology A, 178, 699–709. 10.1007/BF00227381

76. Giurfa, M., Zhang, S., Jenett, A., Menzel, R., & Srinivasan, M. V. (2001). The concepts of ‘sameness’ and ‘difference’in an insect. Nature, 410 (6831), 930–933. 10.1038/35073582

77. Gómez-Laplaza, L. M., & Gerlai, R. (2013). Quantification abilities in angelfish (*Pterophyllum scalare*): The influence of continuous variables. Animal Cognition, 16, 373–383. 10.1007/s10071-012-0578-7

78. Grasso, P. A., Anobile, G., Arrighi, R., Burr, D. C., & Cicchini, G. M. (2022). Numerosity perception is tuned to salient environmental features. IScience, 25 (4). 10.1016/j.isci.2022.104104

79. Greenacre, L., Garcia, J. E., Chan, E., Howard, S. R., & Dyer, A. G. (2022). Vertical versus horizontal Spatial-Numerical Associations (SNA): A processing advantage for the vertical dimension. Plos one, 17 (8), e0262559. 10.1371/journal.pone.0262559

80. Gross, H. J., Pahl, M., Si, A., Zhu, H., Tautz, J., & Zhang, S. (2009). Number-based visual generalisation in the honeybee. PLOS ONE, 4 (1), e4263. 10.1371/journal.pone.0004263

81. Guerra, S., Castiello, U., Simonetti, V., Bonato, B., & McCrink, K. (2025). Asymmetrical distribution of supports affect pea plants movement and shape: Evidence of quantity discrimination? PLoS One, 20 (5), e0322859. 10.1371/journal.pone.0322859

82. Guillaume, M., Schiltz, C., & Van Rinsveld, A. (2020). NASCO: A new method and program to generate dot arrays for non-symbolic number comparison tasks. Journal of Numerical Cognition, 6 (1), 129–147. 10.5964/jnc.v6i1.231

83. Guttman, N., & Kalish, H. I. (1956). Discriminability and stimulus generalization. Journal of experimental psychology, 51 (1), 79. 10.1037/h0046219

84. Hauser, M. D., Carey, S., & Hauser, L. B. (2000). Spontaneous number representation in semi-free-ranging rhesus monkeys. Proceedings of the Royal Society of London. Series B: Biological Sciences, 267 (1445), 829–833. 10.1098/rspb.2000.1078

85. Hochman, S., Havedanloo, R., Heysieattalab, S., & Soltanlou, M. (2024, September). How Does Language Modulate the Association Between Number and Space? A Registered Report of a Cross-cultural Study of the SNARC Effect. 10.31234/osf.io/sme3z

86. Howard, S. R., Avarguès-Weber, A., Garcia, J. E., Greentree, A. D., & Dyer, A. G. (2018). Numerical ordering of zero in honey bees. Science, 360 (6393), 1124–1126. 10.1126/science.aar4975

87. Howard, S. R., Avarguès-Weber, A., Garcia, J. E., Greentree, A. D., & Dyer, A. G. (2019a). Numerical cognition in honeybees enables addition and subtraction. Science Advances, 5 (2), eaav0961. 10.1126/sciadv.aav0961

88. Howard, S. R., Avarguès-Weber, A., Garcia, J. E., Greentree, A. D., & Dyer, A. G. (2019b). Surpassing the subitizing threshold: Appetitive-aversive conditioning improves discrimination of numerosities in honeybees. Journal of Experimental Biology, 222 (19), jeb205658. 10.1242/jeb.205658

89. Howard, S. R., Avarguès-Weber, A., Garcia, J. E., Greentree, A. D., & Dyer, A. G. (2019c). Symbolic representation of numerosity by honeybees (*Apis mellifera*): Matching characters to small quantities. Proceedings of the Royal Society B, 286 (1904), 20190238. 10.1098/rspb.2019.0238

90. Howard, S. R., Greentree, J., Avarguès-Weber, A., Garcia, J. E., Greentree, A. D., & Dyer, A. G. (2022). Numerosity categorization by parity in an insect and simple neural network. Frontiers in Ecology and Evolution, 252. 10.3389/fevo.2022.805385

91. Irie-Sugimoto, N., Kobayashi, T., Sato, T., & Hasegawa, T. (2009). Relative quantity judgment by Asian elephants (*Elephas maximus*). Animal Cognition, 12 (1), 193–199. 10.1007/s10071-008-0185-9

92. Jackson, B. N., Gazes, R. P., & Hampton, R. R. (2025). Spatial representation of magnitude in rhesus macaques: Investigating SNARC effects in quantity and size dimensions. Learning & Behavior, 1–17. 10.3758/s13420-025-00685-0

93. Johnson, B. R. (2010). Division of labor in honeybees: Form, function, and proximate mechanisms. Behavioral ecology and sociobiology, 64 (3), 305–316. 10.1007/s00265-009-0874-7

94. Kern, R., Egelhaaf, M., & Srinivasan, M. V. (1997). Edge detection by landing honeybees: Behavioural analysis and model simulations of the underlying mechanism. Vision research, 37 (15), 2103–2117. 10.1016/S0042-6989(97)00013-8

95. Kheradmand, B., & Nieh, J. C. (2019). The role of landscapes and landmarks in bee navigation: A review. Insects, 10 (10), 342. 10.3390/insects10100342

96. Kilian, A., Yaman, S., von Fersen, L., & Güntürkün, O. (2003). A bottlenose dolphin discriminates visual stimuli differing in numerosity. Animal Learning & Behavior, 31 (2), 133–142.

97. Kreuter, N., Christofzik, N., Niederbremer, C., Bollé, J., & Schluessel, V. (2021). Counting on numbers-numerical abilities in grey bamboo sharks and ocellate river stingrays. Animals, 11 (9), 2634. 10.3390/ani11092634

98. Krusche, P., Uller, C., & Dicke, U. (2010). Quantity discrimination in salamanders. Journal of Experimental Biology, 213 (11), 1822–1828. 10.1242/jeb.039297

99. Kubo, N. (2022). Changes in pigeons’ responses to numerical stimuli depending on total element area differences between stimuli. The Psychological Record, 72 (1), 33–41. 10.1007/s40732-020-00437-8

100. Kuo, J.-C. Z., Ng, L., Stuart-Fox, D., Dyer, A. G., & Howard, S. R. (2025). Spatial preferences influence associations between magnitude and space in honey bees. Animal Behaviour, 221, 123054. 10.1016/j.anbehav.2024.123054

101. Kuzmina, Y., & Malykh, S. (2022). The effect of visual parameters on nonsymbolic numerosity estimation varies depending on the format of stimulus presentation. Journal of Experimental Child Psychology, 224, 105514. 10.1016/j.jecp.2022.105514

102. Lehrer, M., Srinivasan, M. V., & Zhang, S. (1990). Visual edge detection in the honeybee and its chromatic properties. Proceedings of the Royal Society of London. B. Biological Sciences, 238 (1293), 321–330. 10.1098/rspb.1990.0002

103. Lehrer, M., & Srinivasan, M. (1993). Object detection by honeybees: Why do they land on edges? Journal of Comparative Physiology A, 173 (1), 23–32. 10.1007/BF00209615

104. Leibovich, T., Katzin, N., Harel, M., & Henik, A. (2017). From “sense of number” to “sense of magnitude”: The role of continuous magnitudes in numerical cognition. Behavioral and Brain Sciences, 40, e164. 10.1017/S0140525X16000960

105. Lindskog, M., Poom, L., & Winman, A. (2021). Attentional bias induced by stimulus control (ABC) impairs measures of the approximate number system. Attention, Perception, & Psychophysics, 83 (4), 1684–1698. 10.3758/s13414-020-02229-2

106. Lochner, S., Honerkamp, D., Valada, A., & Straw, A. D. (2024). Reinforcement learning as a robotics-inspired framework for insect navigation: From spatial representations to neural implementation. Frontiers in Computational Neuroscience, 18, 1460006. 10.3389/fncom.2024.1460006

107. Loconsole, M., Regolin, L., & Rugani, R. (2023). Asymmetric number-space association leads to more efficient processing of congruent information in domestic chicks. Frontiers in Behavioral Neuroscience, 17, 1115662. 10.3389/fnbeh.2023.1115662

108. Lucon-Xiccato, T., Gatto, E., & Bisazza, A. (2018). Quantity discrimination by treefrogs. Animal behaviour, 139, 61–69. 10.1016/j.anbehav.2018.03.005

109. MaBouDi, H., Barron, A. B., Li, S., Honkanen, M., Loukola, O. J., Peng, F., Li, W., Marshall, J. A., Cope, A., Vasilaki, E., et al. (2021). Non-numerical strategies used by bees to solve numerical cognition tasks. Proceedings of the Royal Society B, 288 (1945), 20202711. 10.1098/rspb.2020.2711

110. McCullagh, P., & Nelder, J. (1989). Binary data. In Generalized linear models (pp. 98–148).

111. Menzel, R., & Giurfa, M. (2001). Cognitive architecture of a mini-brain: The honeybee. Trends in cognitive sciences, 5 (2), 62–71. 10.1016/S1364-6613(00)01601-6

112. Messina, A., Potrich, D., Perrino, M., Sheardown, E., Miletto Petrazzini, M. E., Luu, P., Nadtochiy, A., Truong, T. V., Sovrano, V. A., Fraser, S. E., Brennan, C. H., & Vallortigara, G. (2022). Quantity as a Fish Views It: Behavior and Neurobiology. Frontiers in Neuroanatomy, 16, 943504. 10.3389/fnana.2022.943504

113. Nieder, A. (2021). Neuroethology of number sense across the animal kingdom. Journal of Experimental Biology, 224 (6), jeb218289. 10.1242/jeb.218289

114. Núñez, R. E. (2017). Is there really an evolved capacity for number? Trends in cognitive sciences, 21 (6), 409–424. 10.1016/j.tics.2017.03.005

115. Odic, D., Libertus, M. E., Feigenson, L., & Halberda, J. (2013). Developmental change in the acuity of approximate number and area representations. Developmental psychology, 49 (6), 1103. 10.1037/a0029472

116. Pahl, M., Si, A., & Zhang, S. (2013). Numerical cognition in bees and other insects. Frontiers in psychology, 4, 162. 10.3389/fpsyg.2013.00162

117. Palomero-Gallagher, N., & Amunts, K. (2022). A short review on emotion processing: A lateralized network of neuronal networks. Brain Structure and Function, 227 (2), 673–684. 10.1007/s00429-021-02331-7

118. Petrazzini, M. E. M., & Wynne, C. D. (2016). What counts for dogs (*Canis lupus familiaris*) in a quantity discrimination task? Behavioural processes, 122, 90–97. 10.1016/j.beproc.2015.11.013

119. Piazza, M., Izard, V., Pinel, P., Le Bihan, D., & Dehaene, S. (2004). Tuning curves for approximate numerosity in the human intraparietal sulcus. Neuron, 44 (3), 547–555. 10.1016/j.neuron.2004.10.014

120. Pisa, P. E., & Agrillo, C. (2009). Quantity discrimination in felines: A preliminary investigation of the domestic cat (*Felis silvestris catus*). Journal of Ethology, 27 (2), 289–293. 10.1007/s10164-008-0121-0

121. Pitt, B., Casasanto, D., & Piantadosi, S. T. (2023). No clear evidence for an innate left-to-right mental number line. Proceedings of the National Academy of Sciences, 120 (28), e2306099120. 10.1073/pnas.2306099120

122. Pitt, B., Ferrigno, S., Cantlon, J. F., Casasanto, D., Gibson, E., & Piantadosi, S. T. (2021). Spatial concepts of number, size, and time in an indigenous culture. Science Advances, 7 (33), eabg4141. 10.1126/sciadv.abg4141

123. Pomè, A., Anobile, G., Cicchini, G. M., & Burr, D. C. (2019). Different reaction-times for subitizing, estimation, and texture. Journal of Vision, 19 (6), 14–14. 10.1167/19.6.14

124. Potrich, D., Zanon, M., & Vallortigara, G. (2022). Archerfish number discrimination. Elife, 11, e74057. 10.7554/eLife.74057

125. Price, G. R., Palmer, D., Battista, C., & Ansari, D. (2012). Nonsymbolic numerical magnitude comparison: Reliability and validity of different task variants and outcome measures, and their relationship to arithmetic achievement in adults. Acta psychologica, 140 (1), 50–57. 10.1016/j.actpsy.2012.02.008

126. Regolin, L., Loconsole, M., Rosa-Salva, O., Brosche, K., Macchinizzi, M., Felisatti, A., & Rugani, R. (2025). Numerical cognition in birds. Nature Reviews Psychology, 1–15. 10.1038/s44159-025-00480-8

127. Rigosi, E., Haase, A., Rath, L., Anfora, G., Vallortigara, G., & Szyszka, P. (2015). Asymmetric neural coding revealed by in vivo calcium imaging in the honey bee brain. Proceedings of the Royal Society B: Biological Sciences, 282 (1803), 20142571. 10.1098/rspb.2014.2571

128. Rivas-Blanco, D., Pohl, I.-M., Dale, R., Heberlein, M. T. E., & Range, F. (2020). Wolves and dogs may rely on non-numerical cues in quantity discrimination tasks when given the choice. Frontiers in Psychology, 11, 573317. 10.3389/fpsyg.2020.573317

129. Rodríguez, C., & Ferreira, R. A. (2023). To what extent is dot comparison an appropriate measure of approximate number system? Frontiers in Psychology, 13, 1065600. 10.3389/fpsyg.2022.1065600

130. Rugani, R., Kelly, D. M., Szelest, I., Regolin, L., & Vallortigara, G. (2010). Is it only humans that count from left to right? Biology Letters, 6 (3), 290–292. 10.1098/rsbl.2009.0960

131. Rugani, R., Platt, M. L., Zhang, Y., & Brannon, E. M. (2024). Magnitude shifts spatial attention from left to right in rhesus monkeys as in the human mental number line. Iscience, 27 (2). 10.1016/j.isci.2024.108866

132. Rugani, R., Regolin, L., & Vallortigara, G. (2007). Rudimental numerical competence in 5-day-old domestic chicks (*Gallus gallus*): Identification of ordinal position. Journal of Experimental Psychology: Animal Behavior Processes, 33 (1), 21. 10.1037/0097-7403.33.1.21

133. Rugani, R., Regolin, L., & Vallortigara, G. (2008). Discrimination of small numerosities in young chicks. Journal of experimental psychology: Animal behavior processes, 34 (3), 388. 10.1037/0097-7403.34.3.388

134. Rugani, R., Regolin, L., & Vallortigara, G. (2010). Imprinted numbers: Newborn chicks’ sensitivity to number vs. continuous extent of objects they have been reared with. Developmental science, 13 (5), 790–797.

135. Rugani, R., Vallortigara, G., Priftis, K., & Regolin, L. (2015). Number-space mapping in the newborn chick resembles humans’ mental number line. Science, 347 (6221), 534–536. 10.1126/science.aaa1379

136. Rugani, R., Vallortigara, G., Priftis, K., & Regolin, L. (2020). Numerical magnitude, rather than individual bias, explains spatial numerical association in newborn chicks. eLife, 9, e54662. 10.7554/eLife.54662

137. Rugani, R., Vallortigara, G., & Regolin, L. (2014). From small to large: Numerical discrimination by young domestic chicks (*Gallus gallus*). Journal of Comparative Psychology, 128 (2), 163. 10.1037/a0034513

138. Rugani, R., Vallortigara, G., Vallini, B., & Regolin, L. (2011). Asymmetrical number-space mapping in the avian brain. Neurobiology of learning and memory, 95 (3), 231–238. 10.1016/j.nlm.2010.11.012

139. Schaffer, A., Caicoya, A. L., Widdig, A., Holland, R., & Amici, F. (2025). Quantity discrimination in 9 ungulate species: Individuals take item number and size into account to discriminate quantities. Cognition, 254, 105979. 10.1016/j.cognition.2024.105979

140. Seeley, T. D. (1983). Division of labor between scouts and recruits in honeybee foraging. Behavioral ecology and sociobiology, 12 (3), 253–259. 10.1007/BF00290778

141. Seeley, T. D. (2009). The wisdom of the hive: The social physiology of honey bee colonies. Harvard University Press.

142. Shaki, S., & Fischer, M. H. (2008). Reading space into numbers-a cross-linguistic comparison of the SNARC effect. Cognition, 108 (2), 590–599. 10.1016/j.cognition.2008.04.001

143. Shaki, S., & Fischer, M. H. (2020). Nothing to dance about: Unclear evidence for symbolic representations and numerical competence in honeybees. A Comment on: Symbolic representation of numerosity by honeybees (Apis mellifera): Matching characters to small quantities. Proceedings of the Royal Society B, 287 (1925), 20192840. 10.1098/rspb.2019.2840

144. Shilat, Y., Henik, A., Galili, H., Wasserman, S., Salzmann, A., & Salti, M. (2024). A methodological framework for stimuli control: Insights from numerical cognition. Advances in Methods and Practices in Psychological Science, 7 (4), 25152459241249185. 10.1177/25152459241249185

145. Shilat, Y., Salti, M., & Henik, A. (2021). Shaping the way from the unknown to the known: The role of convex hull shape in numerical comparisons. Cognition, 217, 104893. 10.1016/j.cognition.2021.104893

146. Skorupski, P., MaBouDi, H., Galpayage Dona, H. S., & Chittka, L. (2018). Counting insects. Philosophical Transactions of the Royal Society B: Biological Sciences, 373 (1740), 20160513. 10.1098/rstb.2016.0513

147. Smets, K., Sasanguie, D., Szuecs, D., & Reynvoet, B. (2015). The effect of different methods to construct non-symbolic stimuli in numerosity estimation and comparison. Journal of Cognitive Psychology, 27 (3), 310–325. 10.1080/20445911.2014.996568

148. Snyder, R. J., Barrett, L. P., Emory, R. A., & Perdue, B. M. (2021). Performance of Asian elephants (*Elephas maximus*) on a quantity discrimination task is similar to that of African savanna elephants (*Loxodonta africana*). Animal Cognition, 24 (5), 1121–1131. 10.1007/s10071-021-01504-5

149. Srinivasan, M., & Lehrer, M. (1988). Spatial acuity of honeybee vision and its spectral properties. Journal of Comparative Physiology A, 162 (2), 159–172. 10.1007/BF00606081

150. Stancher, G., Rugani, R., Regolin, L., & Vallortigara, G. (2015). Numerical discrimination by frogs (Bombina orientalis). Animal Cognition, 18 (1), 219–229. 10.1007/s10071-014-0791-7

151. Starr, A., DeWind, N. K., & Brannon, E. M. (2017). The contributions of numerical acuity and non-numerical stimulus features to the development of the number sense and symbolic math achievement. Cognition, 168, 222–233. 10.1016/j.cognition.2017.07.004

152. Stevens, S. S. (1936). A scale for the measurement of a psychological magnitude: Loudness. Psychological Review, 43 (5), 405. 10.1037/h0058773

153. Szűcs, D., Nobes, A., Devine, A., Gabriel, F. C., & Gebuis, T. (2013). Visual stimulus parameters seriously compromise the measurement of approximate number system acuity and comparative effects between adults and children. Frontiers in psychology, 4, 444. 10.3389/fpsyg.2013.00444

154. Tomonaga, M. (2008). Relative numerosity discrimination by chimpanzees (Pan troglodytes): Evidence for approximate numerical representations. Animal cognition, 11 (1), 43–57. 10.1007/s10071-007-0089-0

155. Towne, W. F., Ritrovato, A. E., Esposto, A., & Brown, D. F. (2017). Honeybees use the skyline in orientation. Journal of Experimental Biology, 220 (13), 2476–2485. 10.1242/jeb.160002

156. Triki, Z., & Bshary, R. (2018). Cleaner fish *Labroides dimidiatus* discriminate numbers but fail a mental number line test. Animal Cognition, 21 (1), 99–107. 10.1007/s10071-017-1143-1

157. Uller, C., & Lewis, J. (2009). Horses (*Equus caballus*) select the greater of two quantities in small numerical contrasts. Animal cognition, 12, 733–738. 10.1007/s10071-009-0225-0

158. Vallortigara, G. (2018). Comparative cognition of number and space: The case of geometry and of the mental number line. Philosophical Transactions of the Royal Society B: Biological Sciences, 373 (1740), 20170120. 10.1098/rstb.2017.0120

159. Vogel, J. J., Bowers, C. A., & Vogel, D. S. (2003). Cerebral lateralization of spatial abilities: A meta-analysis. Brain and cognition, 52 (2), 197–204. 10.1016/S0278-2626(03)00056-3

160. Vonk, J., Torgerson-White, L., McGuire, M., Thueme, M., Thomas, J., & Beran, M. J. (2014). Quantity estimation and comparison in western lowland gorillas (*Gorilla gorilla gorilla*). Animal cognition, 17 (3), 755–765. 10.1007/s10071-013-0707-y

161. Walsh, V. (2003). A theory of magnitude: Common cortical metrics of time, space and quantity. Trends in cognitive sciences, 7 (11), 483–488. 10.1016/j.tics.2003.09.002

162. Walsh, V. (2014). A theory of magnitude: The parts that sum to number. 10.1093/oxfordhb/9780199642342.013.64

163. Wason, P. C. (1960). On the failure to eliminate hypotheses in a conceptual task. Quarterly Journal of Experimental Psychology, 12 (3), 129–140. 10.1080/17470216008416717

164. Wehner, R. (1967). Pattern recognition in bees. Nature, 215 (5107), 1244–1248. 10.1038/2151244a0

165. Wrona, F. J., & Dixon, R. J. (1991). Group size and predation risk: A field analysis of encounter and dilution effects. The American Naturalist, 137 (2), 186–201. 10.1086/285153

166. Yang, T.-I., & Chiao, C.-C. (2016). Number sense and state-dependent valuation in cuttlefish. Proceedings of the Royal Society B: Biological Sciences, 283 (1837), 20161379. 10.1098/rspb.2016.1379

167. Zanon, M., Fraser, S. E., & Vallortigara, G. (2025). Numerical discrimination in danionella. iScience. 10.1016/j.isci.2025.113667

168. Zanon, M., Potrich, D., Bortot, M., & Vallortigara, G. (2022). Towards a standardization of non-symbolic numerical experiments: GeNEsIS, a flexible and user-friendly tool to generate controlled stimuli. Behavior Research Methods, 54 (1), 146–157. 10.3758/s13428-021-01580-y

169. Zebian, S. (2005). Linkages between number concepts, spatial thinking, and directionality of writing: The SNARC effect and the reverse SNARC effect in English and Arabic monoliterates, biliterates, and illiterate Arabic speakers. Journal of Cognition and Culture, 5 (1-2), 165–190. 10.1163/1568537054068660

170. Zohar-Shai, B., Tzelgov, J., Karni, A., & Rubinsten, O. (2017). It does exist! A left-to-right spatial-numerical association of response codes (SNARC) effect among native Hebrew speakers. Journal of Experimental Psychology: Human Perception and Performance, 43 (4), 719–728. 10.1037/xhp0000336

171. Zorzi, M., Priftis, K., & Umiltà, C. (2002). Neglect disrupts the mental number line. Nature, 417 (6885), 138–139. 10.1038/417138a

